# Direct electrochemical cortisol detection via transition-resolved interrogation at a defect-engineered graphene interface

**DOI:** 10.64898/2026.09.13.751286

**Authors:** Alam Mahmud, Chuanzhen Zhao, Chibuike Uwakwe, Kuang-Jung Hsu, Il Rok Choi, Tianyang Chen, Brian C. DeFelice, Ira J. Gray, Jihyun (Luna) Hwang, Dan Ilyn, Thomas W. Redvanly, Laura Rijns, Shiyuan Wei, Ines Weber, Diego Uruchurtu Patino, Yangju Lin, Weilai Yu, Hao Lyu, Yuelang Chen, Chengyi Xu, Baiyu Shi, Andrea Sedano, Jeffrey Heo, Joseph M. DeSimone, Zhenan Bao

## Abstract

Continuous tracking of cortisol is central to understanding human stress physiology, yet direct electrochemical detection without biological receptors remains challenging despite its promise for stable and dynamic sensing. Electrochemical reduction of cortisol typically occurs at highly negative potentials, where parasitic interfacial currents, hydrogen evolution, and substantial capacitive background overlap with the cortisol reduction signal, preventing accurate quantification. We overcome this barrier through an integrated material–measurement strategy combining a defect-engineered graphene interface with a transition-resolved interrogation (TRI) measurement strategy. First, we engineered polybenzimidazole-derived laser-induced graphene containing nitrogen-rich defects while suppressing oxygen-derived functionalities which reduced parasitic background currents within the same cathodic potential regime. Next, we designed TRI to leverage differences in the time-dependent evolution of overlapping cathodic processes to isolate a localized cortisol-associated electrochemical reduction transition. Derivative-domain projection coupled with background estimation enables its reliable quantification. This integrated sensing architecture enables sensitive, selective, and dynamic cortisol detection in both artificial and biological interstitial fluids at low nanomolar concentrations. The response remains reproducible across physiologically relevant variations in pH, ionic strength, temperature, and repeated cycling. Together, these capabilities provide a basis for continuous electrochemical cortisol monitoring for future study of stress physiology.

## Introduction

Cortisol regulates the body’s adaptive stress response across timescales ranging from acute fluctuations to ultradian and circadian rhythms (1–3). Persistent dysregulation has been associated with metabolic dysfunction, cardiovascular disease, immune dysfunction, neurobehavioral impairment, and altered circadian timing (1,4,5). Since these dynamics unfold over time, episodic measurements provide only snapshots of cortisol physiology and can miss transient and rhythmic changes. These factors have motivated technologies capable of continuous, minimally invasive, real-time cortisol monitoring (6,7).

A broad range of cortisol sensing strategies has been previously explored, including immunoassays, optical reporters, aptamer-based field-effect transistors, and electrochemical affinity sensors (8–10). Immunoassays offer high analytical accuracy but require additional labeling procedures and are not suitable for real-time continuous measurements (11,12). Optical and fluorescent systems can achieve high sensitivity but require labeling and remain difficult to miniaturize into robust wearable formats (13,14). Electrochemical transduction is particularly attractive for wearable and implantable applications because it supports compact, low-power instrumentation and straightforward integration into flexible, skin-interfaced systems, enabling increasingly sophisticated platforms for physiological monitoring (15–21).

Specifically for cortisol electrochemical sensing, competitive immunosensors, aptamer-based transistor and electrochemical sensors, molecularly imprinted polymer sensors, and membrane-based bioelectronic devices have been investigated for wearable and non-invasive monitoring (22–28). These approaches have enabled substantial advances in molecular selectivity, device integration, and physiological sampling. However, their performance relies on engineered molecular recognition mechanisms that are susceptible to bioreceptor degradation, signal drift, consequent recalibration requirements, and limited reversibility under continuous operation. Importantly, the high-affinity interactions that enable sensitive detection can compromise dynamic tracking, a central challenge in the development of continuous molecular sensing technologies (29–31). Direct electrochemical reduction of cortisol as a sensing method therefore remains attractive because it does not require a separate molecular recognition layer and instead enables sensing through cortisol’s intrinsic electrochemical activity, offering a potential path toward continuous and dynamically responsive monitoring.

However, selective direct electrochemical detection of cortisol under physiologically relevant conditions remains challenging because cortisol lacks a well-resolved redox signature. Its reduction emerges only at strongly negative potentials (typically < −1.5 V), where parasitic interfacial reduction currents arise from electroactive oxygen-containing surface functionalities, hydrogen evolution, and pronounced charging currents associated with extreme cathodic polarization. The above issues obscure the cortisol-linked reduction feature and prevent reliable quantification (32–40). While some studies have reported direct electrochemical reduction signals, the responses often were poorly resolved, cannot be unambiguously assigned to cortisol because of background noises, or have not been demonstrated in physiologically relevant biological fluids (41–46). Accurate resolution of cortisol’s electrochemical transition under physiologically relevant conditions therefore requires improved control over competing electrochemical processes.

Here, we report direct, receptor-free electrochemical detection of cortisol in physiologically relevant interstitial fluids by first developing a method to produce defect-engineered laser-induced graphene interfaces enriched in nitrogen-containing defect sites and depleted of oxygen functionalities. This engineering substantially reduces parasitic cathodic background and preserves rapid heterogeneous electron-transfer kinetics, yielding a stable, low-noise electrochemical interface. Furthermore, we introduce an electrochemical measurement approach, termed transition-resolved interrogation (TRI), which leverages kinetic differences among competing cathodic processes to selectively resolve the cortisol reduction transition within the deeply cathodic regime. The co-design of the graphene interface and measurement strategy thus establishes a methodology for continuous, receptor-free electrochemical detection of cortisol.

## Results and discussion

### Direct electrochemical detection of cortisol

Direct electrochemical detection of cortisol requires first characterizing its intrinsic reduction behavior and understanding why conventional interrogation fails under physiologically relevant conditions. We therefore characterized cortisol’s electron-transfer response under conditions that suppress competing cathodic reactions and compared it to behavior in aqueous electrolytes.

To determine whether the cortisol molecule (**Fig. S1**) exhibits measurable electroactivity under well-controlled conditions, we first examined its behavior in an aprotic environment using cyclic voltammetry (CV) and square-wave voltammetry (SWV). In anhydrous acetonitrile, where hydrogen evolution is suppressed and the interfacial background remains stationary (Supplementary Note 1), cortisol exhibits a quasi-reversible reduction at approximately −2.8 V versus Fc/Fc⁺. This appears as a broad cathodic peak with a smaller anodic return wave in cyclic voltammetry (**Fig. S2**) and a corresponding reverse peak in square-wave voltammetry (**Fig. S3**), demonstrating measurable electrochemical reduction in the absence of competing cathodic reactions.

In aqueous media, the same reduction shifts to approximately −1.75 V versus Ag/AgCl but occurs within a cathodic potential regime that overlaps with hydrogen evolution (**Fig. S4** and Supplementary Note 2). Under these conditions, conventional electrochemical interrogation does not cleanly resolve the cortisol response: no anodic return wave appears in CV (**Fig. S4**), no corresponding reverse peak appears in SWV (**Fig. S5**), and the cortisol-linked feature coincides with the onset of hydrogen evolution reaction (HER). Because the cathodic background evolves during the measurement, subtraction-based correction strategies are ineffective, motivating an interrogation strategy that resolves cortisol under non-stationary cathodic conditions.

To overcome this limitation, we developed a transition-resolved interrogation (TRI) method, a measurement strategy that uses high-amplitude square-wave excitation with defined temporal segments to separate competing cathodic processes in the time domain based on differences in their response times, despite substantial overlap in potential (**Fig. 1a**). Double-layer charging is expected to relax rapidly, whereas irreversible electron-transfer reactions and hydrogen evolution should persist over longer portions of the applied waveform (47–49). These differences in temporal behavior allow TRI to selectively sample regions of waveforms that enrich the cortisol-linked response while minimizing contributions from competing cathodic processes. TRI was implemented on a nitrogen defect-rich graphene interface formed by laser writing of polybenzimidazole (PBI), producing a porous conductive carbon network with abundant electrochemically active defect sites while maintaining a percolated conductive sp² framework (**Fig. 1b**). The interface was subsequently subjected to chemical reduction at an elevated temperature to suppress oxygen-derived surface functionalities that contribute to parasitic background currents within the cortisol reduction window while preserving the underlying defect-rich carbon framework. This combined materials–measurement strategy enables continuous, label-free cortisol sensing despite continuously evolving background currents (**Fig. 1c**).

**Fig. 1.**
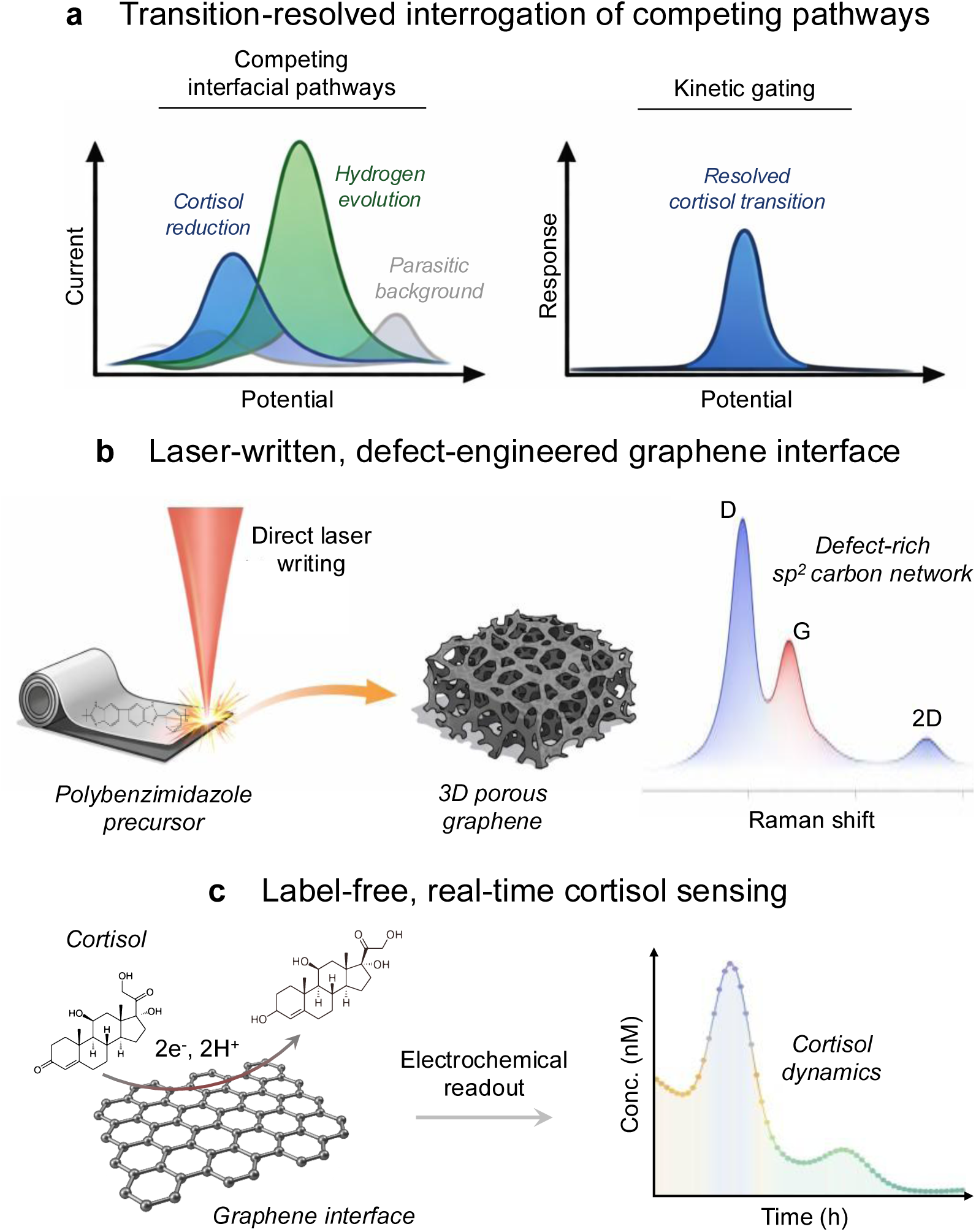
Conceptual framework for direct electrochemical cortisol sensing. **a,** Cortisol electrochemical reduction occurs in a cathodic potential regime where hydrogen evolution, parasitic interfacial reactions, and capacitive background currents overlap with and obscure the cortisol-linked signal. Transition-resolved interrogation (TRI) applies kinetic gating to resolve the cortisol-associated electrochemical feature. **b,** Direct laser writing of polybenzimidazole produces a porous, nitrogen defect-rich graphene network that supports repeated high-amplitude excitation. **c,** The integrated materials–measurement strategy resolves the analyte-linked transition under evolving background conditions, enabling continuous label-free cortisol sensing.

### Transition-resolved interrogation for direct cortisol electroanalysis

TRI builds on the square-wave voltammetric method, whose differential forward–reverse sampling provides high analytical sensitivity and effective discrimination against charging background (50,51). Although conventional SWV can probe both reversible and irreversible electrochemical processes, it is most naturally suited to systems that generate well-defined differential responses under near-equilibrium conditions (48,52). TRI extends these measurement principles beyond conventional background rejection by selectively interrogating temporal regions of the waveform where irreversible electrochemical transitions are strongly expressed relative to competing cathodic processes.

TRI is implemented using high-amplitude square-wave excitation with a cathodic dwell time selected to enhance the cortisol-linked faradaic response relative to competing cathodic processes. Under these conditions, the dwell period extends beyond the rapid decay of capacitive charging while remaining sufficiently short to limit the contribution of hydrogen evolution to the measured response. This creates a kinetic window in which the cortisol-linked reduction transition emerges more distinctly from the surrounding cathodic background. Furthermore, we use a derivative-domain projection, in which the current–potential response is transformed into its first derivative with respect to potential, to sharpen the transition and facilitate quantitative analysis. Together, **Fig. 2a,b** illustrate the conceptual basis of TRI: derivative-domain resolution of the cortisol-linked transition and the excitation-dependent temporal ordering of competing interfacial processes. Additional mechanistic analysis of this excitation-driven regime is provided in Supplementary Note 3.

**Fig 2.**
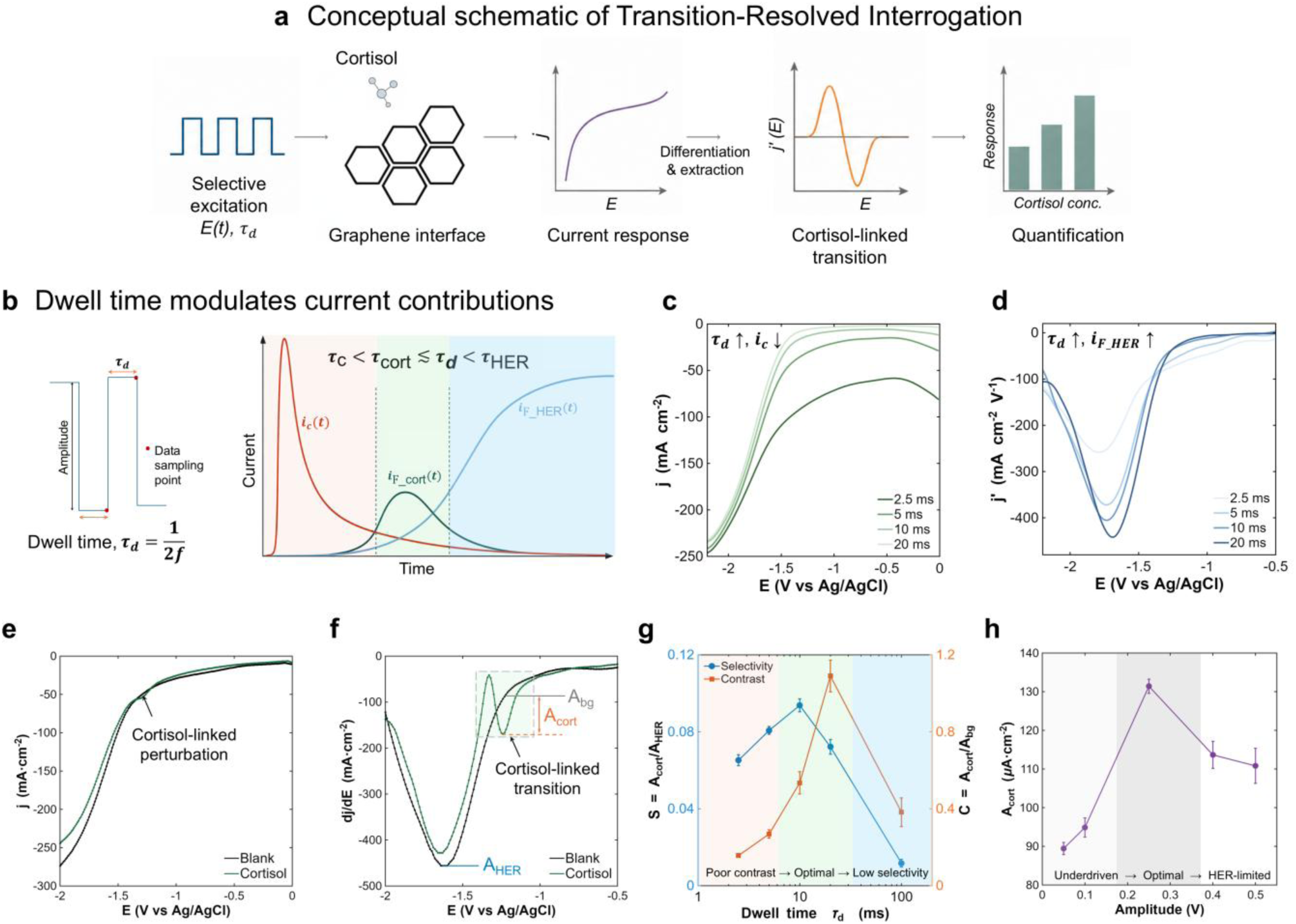
Transition-resolved interrogation (TRI) enables direct electrochemical detection of cortisol in a deep-cathodic regime. **a,** Conceptual schematic of TRI, illustrating high-amplitude excitation with defined dwell time, derivative-domain projection, and baseline estimation to isolate a localized cortisol-linked transition for quantitative readout. **b,** TRI excitation scheme and conceptual timescale ordering of competing current contributions under cathodic polarization, including capacitive charging (*i_c_*), cortisol reduction (*i_F_cort_*), and hydrogen evolution (*i_F_HER_*). **c,d,** Dwell-time-dependent current responses (**c**) and corresponding first-derivative representations (**d**), showing decreasing capacitive and increasing HER contributions with increasing dwell time. **e,** Representative current response under TRI conditions ( *τ_d_* = 10 ms, V= 0.25 V), showing a localized cortisol-linked transition embedded within an evolving cathodic background. **f,** First-derivative representation of the response in **e**, which enhances the identification of the cortisol reduction transition while suppressing slowly varying background contributions. The apparent peak cortisol reduction potential corresponds to the zero-crossing point between the positive and negative lobes of the cortisol-linked derivative feature, whereas *A_cort_* is extracted from the negative lobe relative to the background. **g,** Selectivity (*A_cort_*/*A_HER_*) and contrast (*A_cort_*/*A_bg_*) as a function of excitation dwell time, identifying an intermediate kinetic regime in which cortisol reduction remains observable while competing background processes are constrained. **h,** Dependence of cortisol-transition magnitude on excitation amplitude, showing a non-monotonic response with maximum contrast at intermediate amplitude. Measurements in **g** and **h** were performed using 1000 µM cortisol in 1× PBS containing 0.1 M KCl on reduced PBI-derived laser-induced graphene (PBI-LIG) electrodes. Values are means ± s.d. (n = 3 electrodes). **E**, electrode potential. **j**, current density. **dj/dE**, first-order derivative of the current density with respect to potential. **Ag/AgCl**, silver–silver chloride reference electrode.

We examined these principles experimentally by systematically varying excitation frequency, which directly controls the cathodic dwell time (*τ_d_* = 1/2*f*) and therefore the temporal window available for competing interfacial processes to evolve. The measured current responses show that increasing dwell time allows greater relaxation of capacitive charging (**Fig. 2c**), while the corresponding derivative-domain responses reveal a progressively increasing hydrogen-evolution contribution (**Fig. 2d**). At the longest dwell time tested (20 ms), hydrogen evolution produces a dominant cathodic background that obscures the cortisol-linked transition. As dwell time decreases, hydrogen evolution becomes progressively constrained while the intermediate-timescale cortisol reduction pathway remains accessible. HER-only control measurements further show that hydrogen-evolution currents decrease with increasing excitation frequency (**Fig. S8**), consistent with dwell-time-limited hydrogen-bubble nucleation and growth kinetics (53,54). At very short dwell times (≤5 ms), however, incomplete relaxation of capacitive charging increases non-Faradaic background contributions(**Fig. S9**). These opposing temporal dependencies create an intermediate dwell-time regime in which the cortisol-linked response remains accessible while competing background contributions are constrained. We therefore selected a 10 ms dwell time (50 Hz), which maximized selectivity against HER while maintaining substantial contrast against the background (**Fig. 2g**).

Under the selected TRI conditions, the measured current response exhibits a cortisol-linked transition embedded within the broader cathodic background (**Fig. 2e**). Taking the first derivative of the current response with respect to potential (**Fig. 2f**) enhances the resolution of this transition relative to slowly varying background contributions, improving its discrimination. Such derivative voltammetry has previously been used to resolve overlapping voltammetric features (55,56). The resulting derivative-domain feature enables extraction of the cortisol-linked response for quantitative analysis.

To quantify these competing effects, we defined a selectivity metric, *S*(*τ_d_*) = *A_cort_*/*A_HER_*, where *A_cort_* denotes the amplitude of the cortisol-linked transition and *A_HER_* the hydrogen-evolution contribution, and a contrast metric, *C*(*τ_d_*) = *A_cort_*/*A_bg_*, where *A_bg_* represents the residual background contribution. These complementary metrics capture suppression of hydrogen evolution and capacitive background, respectively. Their trends define an intermediate kinetic regime (∼10–20 ms) in which cortisol reduction remains resolvable while both hydrogen evolution and capacitive distortion are constrained (**Fig. 2g**). The cortisol-linked transition magnitude (*A*_cort_) was subsequently used as the analytical signal for quantitative cortisol measurements throughout the study.

To define how excitation amplitude shapes TRI performance, we systematically varied the square-wave voltage amplitude to probe the trade-off between activating the cortisol reduction pathway and competing background processes. The transition magnitude (*A_cort_*) exhibits a non-monotonic dependence on voltage amplitude (**Fig. 2h**). At low amplitudes, it is insufficient to elicit a pronounced cortisol-linked transition, resulting in a weak analyte response. As amplitude increases, the transition becomes progressively stronger, yielding an enhanced cortisol-linked response.

At higher amplitudes, however, increased capacitive charging and accelerated hydrogen evolution reduce the relative prominence of the analyte signal. This creates an intermediate amplitude regime that maximizes transition contrast. Increasing amplitude also produces a monotonic positive shift in the apparent reduction potential of the cortisol-linked transition (**Fig. S10**), indicating that the reduction process becomes observable earlier within the excitation waveform under stronger cathodic polarization.

### Defect-engineered PBI-LIG interface for sensitive cortisol detection

Graphene-based electrodes provide electrochemical interfaces with wide potential windows, tunable defect landscapes, and rapid heterogeneous electron-transfer (HET) kinetics that are attractive for redox-active small-molecule sensing (57–59). To realize these advantages in a scalable, patterned electrode format, we employed laser-induced graphene (LIG), a direct-write approach that converts polymer precursors into porous conductive carbon (60). While polyimide (PI) is a widely used precursor for laser-induced graphene, polybenzimidazole (PBI) was selected here because its nitrogen-rich, oxygen-free aromatic backbone has potential to yield a heteroatom-doped graphene-like carbon following laser conversion (61–63). A benchmark comparison between PBI-derived and PI-derived LIG is provided in Supplementary Note 4 and **Fig. S6**.

We first characterized the as-written PBI-derived LIG (PBI-LIG) to define the structural features relevant to electrochemical sensing. Raman spectroscopy (**Fig. 3a**) shows pronounced D and G bands at ∼1325 and ∼1580 cm^-1^, an elevated D/G ratio (∼1.9), and a modest 2D band, consistent with a defect-rich graphene-like carbon composed of small, electronically connected sp^2^ domains. The resolved D′ feature indicates mixed edge-and vacancy-type defects characteristic of laser-written carbon.

**Fig. 3.**
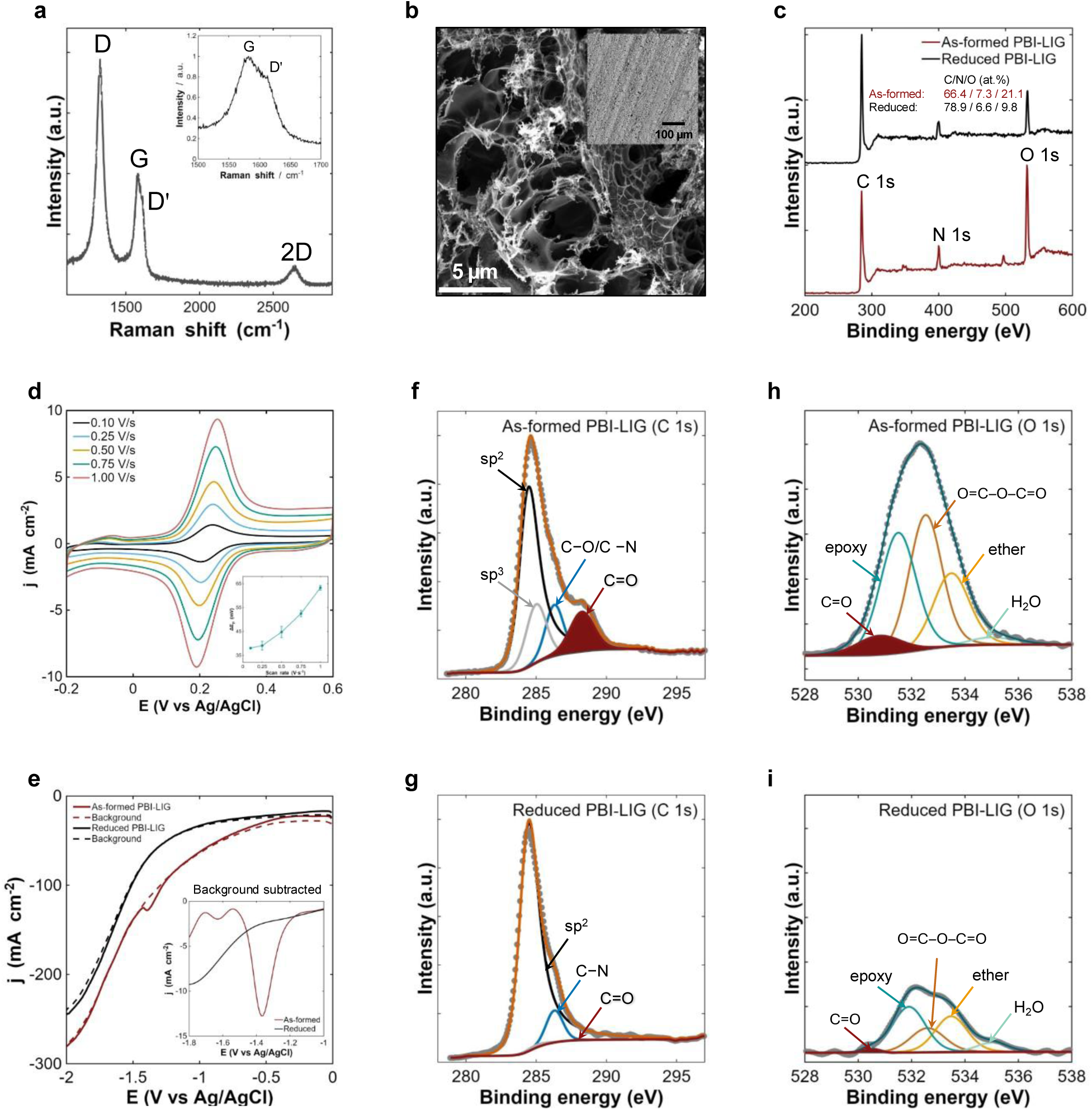
Defect-engineered PBI-derived graphene stabilizes the electrochemical interface for low-noise sensing. **a,** Raman spectrum of PBI-LIG showing a prominent D band, elevated D/G ratio (∼1.9), and modest 2D band, consistent with defect-active graphene-like carbon composed of small, electronically connected sp² domains. Inset: resolved G and D′ features indicating mixed edge- and vacancy-type defects. **b,** Scanning electron microscopy (SEM) image of PBI-LIG showing a porous, interconnected morphology. Scale bar: 5 µm. Inset: lower-magnification view (500 × 500 µm). **c,** XPS survey spectra of as-formed and reduced PBI-LIG. Both spectra confirm a heteroatom-containing carbon framework, with the reduced PBI-LIG exhibiting substantially lower oxygen content while largely preserving nitrogen content. **d,** Cyclic voltammetry of ferri/ferrocyanide across varying scan rates, showing compressed peak-to-peak separations (ΔEₚ) and weak scan-rate dependence, consistent with rapid heterogeneous electron transfer. Inset: ΔEₚ plotted as a function of scan rate. **e,** Raw electrochemical responses of as-formed and reduced PBI-LIG in blank buffer, showing suppression of the parasitic cathodic background following reduction. Following conditioning in buffer (see Methods and **Fig. S18**), an electrochemical response was recorded and used as the background. The electrodes were then incubated in blank buffer for 10 min and measured again. Inset: background-subtracted responses highlighting the electrochemical features that develop during the 10-min blank incubation. **f,g,** High-resolution XPS C 1s spectra of as-formed and reduced PBI-LIG. **h,i,** XPS O 1s spectra of as-formed and reduced PBI-LIG, confirming reduction of oxygen-containing surface functionalities. For comparison of relative oxygen content, the O 1s spectra were normalized to the maximum C 1s intensity of the corresponding sample.

Scanning electron microscopy (**Fig. 3b**) reveals a porous, interconnected morphology that provides high surface area and distributed electron-transfer pathways. Complementary XPS survey spectra (**Fig. 3c**) confirm a heteroatom-containing carbon framework comprising substantial nitrogen incorporation and oxygen-containing surface functionalities.

Consistent with this defect-rich sp^2^ framework, PBI-LIG supports rapid and reproducible HET. Ferri/ferrocyanide voltammetry (**Fig. 3d, Fig. S11**) shows compressed peak-to-peak separations (ΔEₚ ≈ 38–65 mV across 0.1–1.0 V s^-1^) with weak scan-rate dependence, reflecting fast electron exchange across the interconnected, non-planar carbon network (64–66). These results establish the electrochemical suitability and reproducibility of PBI-LIG, supporting its use as a suitable platform for direct cortisol interrogation.

Laser conversion in ambient conditions introduces oxygen-based functionalities—most prominently carbonyl, epoxide, and hydroxyl groups—that undergo intrinsic electroreduction and generate parasitic faradaic currents in analyte-free electrolyte (34, 57–59,67). High-resolution XPS C 1s and O 1s spectra (**Fig. 3f–i**) confirm these oxygen-linked species; C 1s spectra (**Fig. 3f**) confirm substantial carbonyl, ether/amine, and other oxygen-derived components, while the corresponding O 1s spectrum (**Fig. 3g**) shows strong oxygen-derived contributions. Electrochemical interrogation of as-formed PBI-LIG (**Fig. 3e**) accordingly reveals a pronounced background feature in blank buffer within the same potential window as cortisol reduction.

To eliminate this interference, we applied sequential hydrazine reduction and thermal annealing to remove oxygen-based defects and partially restore the sp² network. Hydrazine reduction is used to reduce carbonyl and epoxide groups, converting them into volatile or weakly bound species without disrupting the conjugated carbon lattice or stripping the nitrogen dopants (68–70). A subsequent mild thermal annealing step removes residual hydrazine-derived fragments and weakly bound oxygenated species, helps to repair the sp^2^ framework and yields a cleaner, more electronically uniform graphene interface. Consistent with this processing, XPS survey spectra (**Fig. 3c**) show oxygen removal while largely preserving the nitrogen content.. High-resolution XPS further confirms this transformation. The reduced PBI-LIG C 1s spectrum (**Fig. 3h**) shows near-complete suppression of carbonyl and epoxide components, while the C 1s-normalized O 1s spectrum (**Fig. 3i**) exhibits a substantial reduction in oxygen-linked features.

The electrochemical consequences are shown in **Fig. 3e** and **Fig. S11**. Reduced PBI-LIG exhibits markedly lower parasitic faradaic background currents, yielding a clean and stable baseline for subsequent cortisol sensing. Blank-buffer electrochemical characterization further reveals broader changes in the interfacial electrochemical response (SI Note 5, **Fig. S7**). High-resolution N 1s spectra further reveal a redistribution of nitrogen chemical environments, with an increased relative contribution of pyridinic-like nitrogen and a decreased pyrrolic contribution (**Fig. S7c,d**). Pyridinic nitrogen has been associated with proton-coupled electron transfer on carbon surfaces in previous studies (71). Together, these observations indicate that the reduced PBI-LIG provides a defect-engineered graphene interface capable of supporting selective cortisol detection.

### Cortisol sensing performance

We evaluated quantitative cortisol sensing in a physiologically relevant buffer (1× PBS + 0.1 M KCl) using TRI and the defect-engineered graphene interface. Measurements were performed in a three-electrode configuration (**Fig. 4a**) to ensure stable potential control and minimize electrode polarization throughout interrogation.

**Fig. 4.**
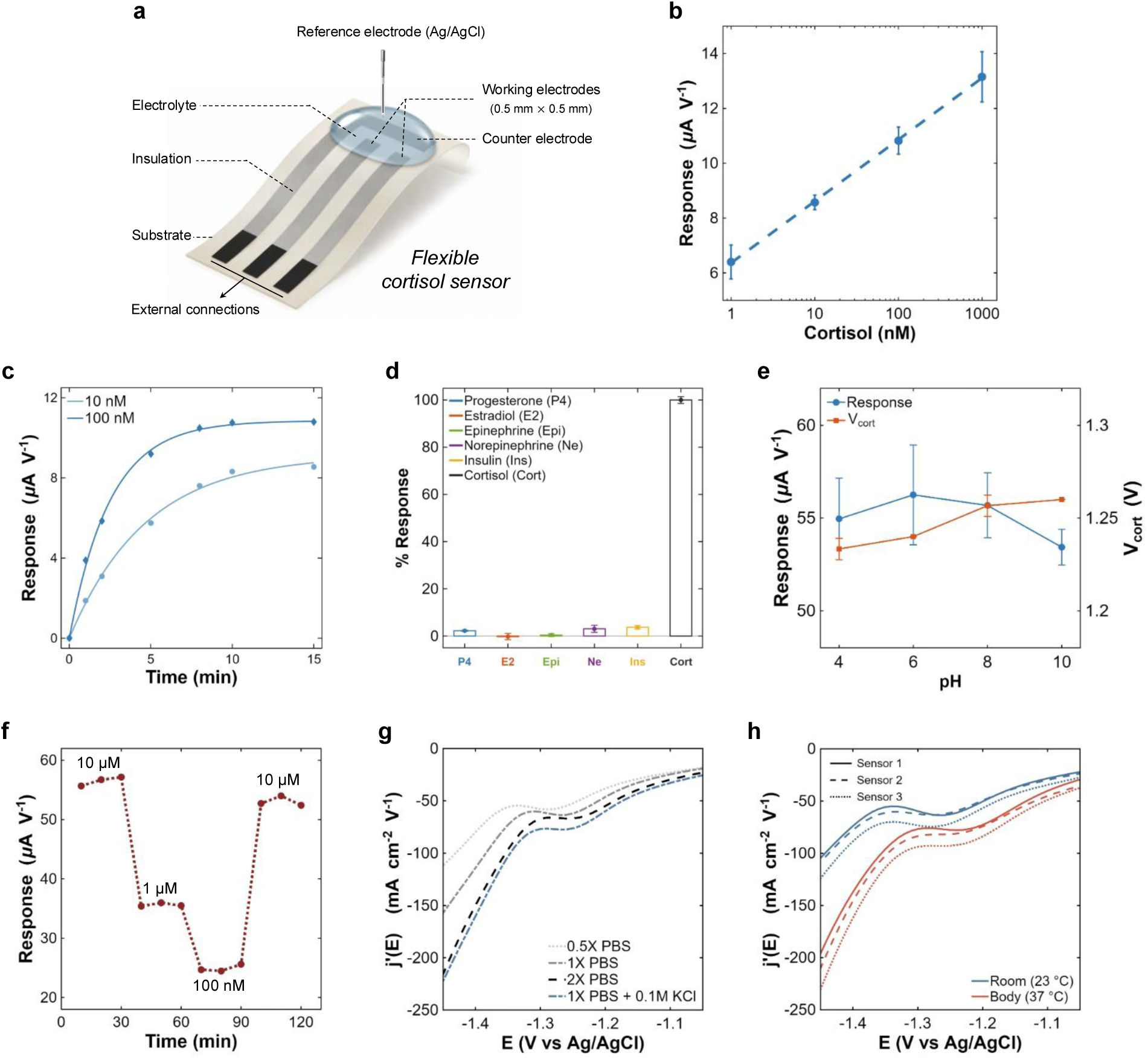
Quantitative behavior of the sensing interface under dynamic interrogation. **a,** Schematic of the flexible three-electrode cortisol sensor and measurement configuration, showing the reduced PBI-LIG working and counter electrodes, Ag/AgCl reference electrode, electrolyte droplet, and key device dimension. **b,** Transition-resolved responses increase linearly with logarithm of cortisol concentration, yielding a well-defined calibration that supports quantitative readout (n= 4). **c,** Time-dependent cortisol responses at 10 and 100 nM approach steady state within ∼10 min, compatible with the minutes-scale dynamics of physiological cortisol fluctuations (also see **Fig. S13**) (n = 1 per concentration). **d,** Structurally related steroids and representative interferents generate negligible transitions under identical interrogation conditions confirming chemical specificity (also see Fig S13) (n=3). 100 nM cortisol (Cort) and representative physiological interferents (200 pM progesterone (P4), 10 nM estradiol (E2), 10 nM epinephrine (Epi), 10 nM norepinephrine (Ne), and 1 nM insulin (Ins). **e,** The apparent reduction potential (∼ −1.24 V) and cortisol-linked response remain stable across physiologically relevant pH values, indicating robustness to acid–base variation (n=4). **f,** Responses from the cortisol-linked transition over consecutive measurement at different cortisol concentrations, demonstrating ability of the defect-engineered graphene interface to detect different concentrations. **g,** Transition characteristics are preserved across distinct ionic environments, supporting operation in diverse physiological media. **h,** The cortisol-linked transition shows modest, systematic thermal shifts between laboratory and physiological temperatures across three sensors, with only minor changes in peak position. Experiment in **g** was independently repeated using three sensors, yielding consistent trends across independent devices.

Under these conditions, the measured response was linear with the logarithm of cortisol concentration over 1–1000 nM (**Fig. 4b**). Time-dependent measurements showed that the signal approached steady state over ∼10 min (**Fig. 4c**), indicating that the response is influenced by interfacial equilibration rather than by bulk concentration alone. This behavior is consistent with contributions from cortisol adsorption and transport at the graphene–electrolyte interface, which influence the local population of electroactive cortisol available for reduction. Adsorption-controlled cortisol electroreduction has previously been reported at carbon microelectrode interfaces (41), supporting surface interactions as a plausible contributor to the response observed here. The analytical sensitivity was determined from the slope of the calibration curve, and the limit of detection (LOD) was estimated to be approximately 0.1 nM using the 3σ criterion based on 4 replications of blank measurements. Responses to structurally related steroids and representative hormonal potential interferents were minimal under identical conditions (**Fig. 4d**).

The sensing response did not change substantially in pH range of 4–10, with both response magnitude and reduction potential largely preserved (**Fig. 4e**). Repeated interrogation over approximately 2 h tracked sequential changes in cortisol concentration while maintaining consistent electrochemical responses (**Fig. 4f**), demonstrating sustained sensor functionality over the measurement period. The cortisol-linked feature is likewise preserved across distinct electrolyte compositions and ionic strengths, demonstrating tolerance to variations in electrolyte composition (**Fig. 4g**). Temperature change from room temperature to body temperature of 37 °C produces modest, systematic shift in the response across the physiological range without compromising observability of the transition (**Fig. 4h**).

### Cortisol sensing across interstitial-fluid environments

To evaluate cortisol sensing across physiologically relevant interstitial-fluid environments, we examined sensor performance in artificial and biological ISF. Dermal ISF was collected using a previously reported minimally invasive puncture–out–press (POP) workflow, followed by centrifugation to recover extracted fluid for downstream analysis (**Fig. 5a**) (72). Representative collection-site images immediately after sampling and 4 h later illustrate the limited visible skin disruption associated with the procedure. TRI-based electrochemical sensing results of cortisol are compared with ground truth from LC–MS/MS quantification **Fig. 5b**. The experimental three-electrode configuration used for electrochemical measurements is shown in **Fig. S14**.

**Fig. 5.**
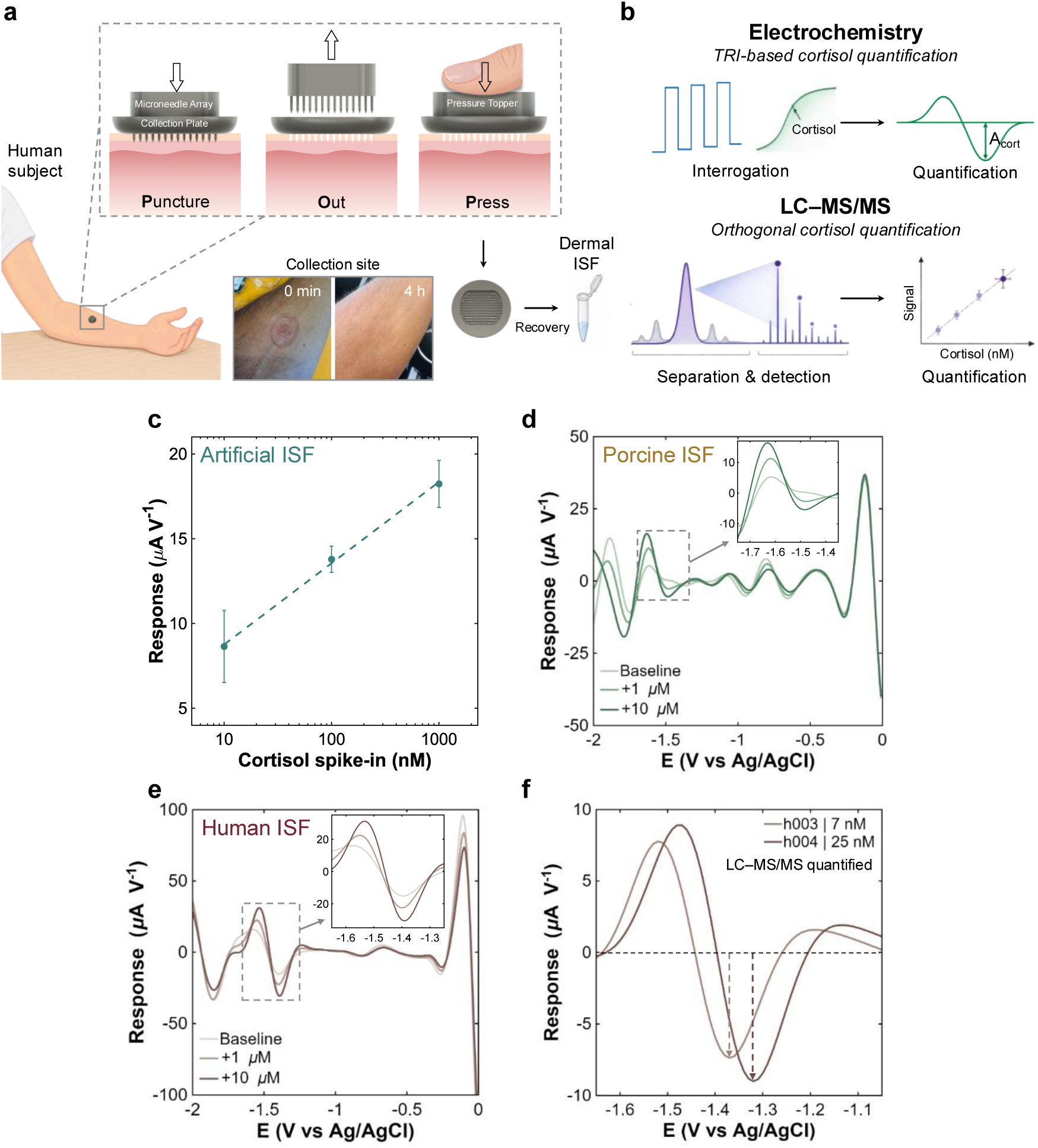
Cortisol detection in artificial and biological interstitial-fluid (ISF). **a,** Dermal ISF sampling workflow using the Puncture–Out–Press (POP) device (72), followed by centrifugation to recover extracted ISF. Representative collection-site images at 0 and 4 h illustrate the limited visible skin disruption associated with the procedure. **b**, Conceptual representation of TRI-based electrochemical and orthogonal LC–MS/MS quantification of cortisol. **c**, Calibration of the cortisol response in artificial ISF over 10–1000 nM, showing a concentration-dependent increase in sensor response. Data represent mean ± s.d. (n = 3). **d,** Representative extracted TRI signatures measured in porcine ISF before and after addition of 1 and 10 µM cortisol. Inset: expanded view of the cortisol-linked transition region. **e,** Representative extracted TRI signatures measured in native human ISF before and after addition of 1 and 10 µM cortisol. Inset: expanded view of the cortisol-linked transition region. For **d** and **e**, measurements were performed using three independently fabricated sensors; representative data from one sensor are shown. Corresponding raw *j–V* curves are provided in **Figs. S15 and S16**, respectively. **f**, Representative extracted TRI signatures from one sensor measured in two native human ISF samples (h003 and h004) containing 7 and 25 nM cortisol, respectively, as independently determined by LC–MS/MS. Measurements from three sensors are shown in **Fig. S17**.

We first evaluated quantitative performance in commercially available artificial interstitial fluid (aISF; Simulated Interstitial Fluid, BZ254, Biochemazone). The extracted response increased monotonically with cortisol concentration over 10–1000 nM, demonstrating quantitative sensing across the physiologically relevant nanomolar range in an interstitial-fluid-like media (**Fig. 5c**). We next examined whether the cortisol-linked electrochemical transition remained identifiable in biological ISF. In porcine ISF, extracted TRI signatures revealed the characteristic cortisol-linked transition following addition of 1 and 10 µM cortisol (**Fig. 5d**; corresponding raw *j–V* curves in **Fig. S15**). We then evaluated native human dermal ISF, where the same additions revealed the transition within the broader background (**Fig. 5e**; corresponding raw *j–V* curves in **Fig. S16**). These higher-concentration additions enabled confident assignment of the cortisol-linked electrochemical transition in complex biological ISF.

Finally, to assess whether the sensor could detect endogenous cortisol at physiological concentrations, we tested it on native human ISF samples. Representative extracted TRI signatures show the cortisol-linked transition in both samples (**Fig. 5f**), with replicate measurements from three independent sensors provided in **Fig. S17**. Cortisol concentrations in h003 and h004, collected from two different human subjects, were independently determined by LC–MS/MS to be 7 and 25 nM, respectively (see Methods). The higher concentration in h004 was accompanied by a larger observed sensor response magnitude. The slight difference in apparent cortisol reduction potential between the samples from two different human subjects may reflect differences in their composition, including pH (**Fig. 4e**), ionic strength (**Fig. 4g**), or other matrix constituents.

Together, these results show that the defect-engineered graphene interface, coupled with TRI, preserves the cortisol-linked electrochemical signature across increasingly complex matrices, spanning quantitative calibration in aISF, transition identification in porcine ISF, and endogenous cortisol detection in native human ISF.

## Conclusion

In this work, we developed a transition-resolved interrogation (TRI) strategy to address the challenge of direct electrochemical cortisol detection in a deeply cathodic regime dominated by hydrogen evolution, capacitive charging, and evolving interfacial background currents. To enable sensitive and selective cortisol detection, we engineered a polybenzimidazole-derived laser-induced graphene interface and applied hydrazine chemical reduction to reduce the amount of oxygen-derived defect states associated with parasitic cathodic background processes. Together, the defect-engineered interface and TRI resolve the cortisol-linked reduction transition under non-stationary cathodic conditions, enabling direct electrochemical readout of cortisol in a deeply cathodic regime that has remained inaccessible to conventional electroanalytical methods.

With further translational development, this approach may support continuous electrochemical monitoring of stress-related physiology. More broadly, these results demonstrate that weak electrochemical signatures can become accessible through co-engineering of the electrochemical interface and interrogation strategy, providing a general route for direct electronic access to challenging molecular targets.

## Methods

### Materials

Polybenzimidazole (PBI) membranes (35 µm thickness, 200 × 200 mm sheets) were purchased from Yatokess (Hong Kong). Hydrazine monohydrate (≥98%) was obtained from TCI Chemicals (Tokyo Chemical Industry, USA). Styrene-butadiene-styrene (SBS) triblock copolymer, toluene (anhydrous, ≥99.8%), acetone (ACS grade), and isopropyl alcohol (IPA, ACS grade) were purchased from Sigma-Aldrich. Polydimethylsiloxane (PDMS; Sylgard 184) was obtained from Dow Corning. Commercial glassy carbon electrodes (GCEs, 3 mm diameter) were purchased from CH Instruments (USA). Ag/AgCl reference electrodes (3 M NaCl; MF-2052) were obtained from BASi (Bioanalytical Systems, Inc., USA). Cortisol (hydrocortisone), progesterone, estradiol, epinephrine, insulin, and norepinephrine were purchased from Sigma-Aldrich. Cortisol-9,11,12,12-*d*_4_, used as the internal standard for LC–MS/MS analysis, was purchased from CDN Isotopes (LGC Standards; catalog no. CDN-D-5280). Anhydrous acetonitrile (MeCN, ≥99.9%) and tetrabutylammonium hexafluorophosphate (TBAPF₆, electrochemical grade) were obtained from Sigma-Aldrich and used to prepare 0.1 M supporting electrolyte for nonaqueous measurements. Potassium ferricyanide (K_3_[Fe(CN)_6_]) and potassium ferrocyanide trihydrate (K_4_[Fe(CN)_6_]·3H_2_O) were purchased from Sigma-Aldrich. Phosphate-buffered saline (PBS, 1×) was obtained from Thermo Fisher Scientific. Artificial interstitial fluid (aISF) was purchased from Biochemazone (Canada). All other solvents and salts were purchased from Sigma-Aldrich or Thermo Fisher Scientific.

### Laser-induced graphene (LIG) formation from PBI

Polybenzimidazole (PBI) membranes (35 μm thickness) were cleaned and thermally pretreated before laser processing to remove processing contaminants and reduce variability associated with moisture content. The films were sequentially cleaned with acetone and isopropyl alcohol, rinsed with fresh isopropyl alcohol, and annealed under vacuum at 235 °C.

The pretreated PBI films were supported on PDMS-coated glass substrates during laser writing. Laser-induced graphene (LIG) patterns were generated using a CO₂ laser cutter (Epilog Fusion M2, 10.6 μm wavelength) operated in raster mode under a localized nitrogen environment. Laser-processing conditions were optimized to produce continuous, mechanically stable PBI-LIG patterns suitable for subsequent electrochemical measurements. Following laser writing, the patterned films were stored under reduced pressure until further processing or characterization.

For comparison, LIG was also formed on commercial polyimide (PI) membranes using corresponding processing conditions. A benchmarking comparison between PBI-LIG and PI-LIG is provided in Supplementary Note 4.

### Chemical modification of PBI-LIG

Patterned PBI-LIG electrodes were treated with hydrazine monohydrate at elevated temperature under nitrogen, thoroughly rinsed with Type II deionized water, and dried under N₂. The chemically treated electrodes were subsequently annealed under reduced pressure before electrochemical characterization and sensing measurements.

### General characterizations

Raman spectra were collected using a Horiba XploRA system equipped with a 638 nm excitation laser and a Syncerity CCD detector (−60 °C). Spectra were acquired using a 100× objective, a 1800 gr/mm grating, a 100 µm entrance slit, and a 300 µm confocal hole. Each spectrum was recorded over the 1000–3090 cm⁻¹ range with a 30 s acquisition time and three accumulations.

X-ray photoelectron spectroscopy (XPS) was performed using a PHI VersaProbe 3 system equipped with a monochromated Al Kα X-ray source (1486.6 eV). Samples were electrically grounded to the sample stage inside the ultrahigh-vacuum chamber. Survey spectra were collected at a pass energy of 224 eV, and high-resolution C 1s and N 1s spectra were acquired at a pass energy of 112 eV. Binding energies were calibrated to the C 1s peak at 284.8 eV, and data were processed and peak-fit using CasaXPS.

Scanning electron microscopy (SEM) images were acquired using an FEI Magellan 400 XHR system operated at 5 kV and 100 pA. Samples were mounted on aluminum stubs using conductive carbon tape and imaged under high-vacuum conditions.

Electrochemical measurements were performed using a CH Instruments 700F bipotentiostat in a three-electrode configuration. The patterned PBI-LIG electrode served as both the working and counter electrodes, and a commercial Ag/AgCl (3 M NaCl) electrode was used as the reference unless otherwise noted. The geometric footprint of each LIG working electrode was 0.5 mm × 0.5 mm. Prior to all measurements; electrodes were annealed on a vacuum heater at 80 °C for 20 min.

### Cortisol sensor fabrication

PBI-LIG working and counter electrodes were patterned according to the predefined device geometry and subjected to the same chemical and thermal post-processing. An insulating layer was stencil-printed to define an exposed 0.5 mm × 0.5 mm LIG working-electrode area, and printed silver contacts were used to provide electrical connection to the electrochemical workstation. The counter electrode was designed with a substantially larger geometric area than the working electrode to minimize counter-electrode polarization during TRI measurements. A commercial Ag/AgCl reference electrode (3 M NaCl; MF-2052, BASi) was used for electrochemical measurements.

### Microneedle-assisted dermal ISF extraction

Porcine ear skin was harvested from recently euthanized Yorkshire pigs following methods adapted from Golombek et al. (73). The outer ear was rinsed with Dulbecco’s phosphate-buffered saline (DPBS), shaved with a disposable razor, and full-thickness skin was excised from the underlying cartilage using a scalpel. Any subcutaneous fat was trimmed with a razor. The isolated skin was briefly sterilized by immersion in a povidone-iodine (Betadine)/DPBS mixture, followed by a second rinse in fresh DPBS. The tissue was patted dry with a Kimwipe and stored at −20 °C. For experiments, frozen skin samples were thawed in DPBS at room temperature in a Petri dish. After thawing, excess surface moisture was removed with a Kimwipe, and the tissue was transferred to a disposable dissecting board. A microneedle array patch (MAP), 3D-printed on a prototype Carbon S2 printer using KeySplint Hard resin (Keyprint #4220004), was applied to the skin using a 3D-printed tunable applicator (TAPP) (74). Following insertion, the MAP was removed, and a 3D-printed suction cup was positioned over the puncture site. Vacuum was applied for 5 min to facilitate ISF extraction. The suction cup was then removed, and ISF was collected from the skin surface using a micropipette and transferred into an Eppendorf tube. Samples were immediately stored at −80 °C until analysis.

### Human ISF collection

Dermal interstitial fluid (ISF) collection was conducted at Stanford University under IRB protocol no. 80484. Inclusion criteria include 18 years of age or older. Exclusion criteria include history of skin conditions that predispose the participant to adverse reactions to skin puncture, active infection, open wound, rash, or skin breakdown on the arm being used for MAP application, and known acrylate allergy. All procedures followed aseptic techniques.

ISF was collected from the volar forearm using a device and protocol adapted from Hung et al (72). A modified version of the Version C (clinical) puncture-out-and-press (POP) device was used. The collection plate and topper were pre-weighted, and a warm pack (HotHands Hand Warmers, Amazon) was applied to the skin site for 5 minutes, while the MAPs were mated to the collection plate. The skin was then wiped with an alcohol pad, and the assembled MAPs and the plate were placed on the arm within the applicator stand. A commercial spring-loaded applicator, modified with a 25-N spring (PC031-281-7500-MW-0599-CG-N-IN, The Spring Store) was deployed four times from predefined positions on the stand to ensure uniform impact across the MAP surface.

The MAPs were then carefully removed while the collection plate remained in place. A topper was added, and a thin-film force sensor (Qinlorgo, RP-C-MK01X, Amazon) placed on top was used to guide application of 750 g of thumb pressure for 5 minutes. The plate and topper were detached from the skin and weighed to quantify the collected ISF. The puncture site was photographed, cleaned with an alcohol pad, and covered with an adhesive bandage.

After ISF collection, the collection plate and topper were gently detached, and 100 µL of DPBS was pipetted onto the topper. The plate was then reattached, and the assembly was placed in a custom adapter on a 15-mL conical tube and centrifuged in 2000 g for 10 seconds. ISF was extracted into the tube, while RBCs settled at the bottom. The upper and lower halves were pipetted into separate Eppendorf tubes: the supernatant fraction (upper half), contaminated only by hemolyzed RBCs, and the sediment fraction (lower half), containing both hemolyzed and intact RBCs. Hung et al. reported such physical decontamination via centrifugation resulted in <1% blood contamination. Both ISF supernatant and sediment were stored at −80 °C until analysis.

### LC–MS analysis of cortisol in ISF

ISF samples were spiked with cortisol-d4 internal standard before extraction, followed by addition of five volumes of ice-chilled methanol. Samples were vortexed for 20 s at 3,000 rpm, shaken at −4 °C for 6 min at 1,200 rpm, and centrifuged at room temperature for 2 min at 14,000 rpm. Supernatants were transferred to fresh tubes, evaporated to dryness using a CentriVap at room temperature, and stored at −80 °C until analysis. Dried samples were reconstituted in 10 µL of 3:2 (v/v) LC–MS-grade acetonitrile:water containing 60 ng mL^-1^ CUDA, vortexed, sonicated for 5 min, and centrifuged at 14,000 × *g* for 2 min before transfer to a 384-well plate.

Chromatographic separation was performed using a Thermo Vanquish UPLC equipped with an Ascentis Express RP-Amide column (150 × 2.1 mm, 2.7 µm). Mobile phase A consisted of 10 mM ammonium formate and 0.125% formic acid in water, and mobile phase B contained the same additives in 95:5 (v/v) acetonitrile:water. Separation was performed at a flow rate of 0.4 mL min⁻¹ using the following gradient: 5% B from 0 to 1.0 min, increased linearly to 95% B at 7.0 min, held at 95% B until 8.0 min, returned to 5% B at 9.0 min, and held at 5% B until 10.0 min. A 5 µL sample volume was injected for analysis.

Mass spectra were acquired using a Thermo Q Exactive HF Hybrid Quadrupole-Orbitrap mass spectrometer in positive-ionization mode using full MS–ddMS² acquisition over *m/z* 60–900, with resolutions of 60,000 for MS¹ and 15,000 for MS², a loop count of 4, and a 1.0-Da isolation window. Cortisol was quantified against an authentic cortisol calibration curve using cortisol-d4 as the stable-isotope internal standard. Cortisol-d4 was added to samples, calibration standards, and blanks at the same concentration before sample processing, and quantification was based on the endogenous cortisol/cortisol-d4 signal ratio using a linear calibration fit spanning the measured ISF cortisol concentrations.

### Electrochemical measurements

#### 1. Electrode quality evaluation

Cyclic voltammetry (CV) was performed in an equimolar solution of 2.0 mM potassium ferricyanide and 2.0 mM potassium ferrocyanide prepared in 1× PBS containing 0.1 M KCl. CV scans were collected using a CH Instruments 700F bipotentiostat in a three-electrode configuration, with the patterned PBI-LIG electrode serving as both the working and counter electrodes and a commercial Ag/AgCl electrode as the reference. Measurements were conducted from −0.2 V to 0.6 V at a scan rate of 0.5 V s⁻¹, using a 10-segment sweep and a 10-mV sampling interval. A 2 s quiet time was applied before each scan. The sensitivity setting was 1×10⁻⁵ A V⁻¹.

#### 2. Material and interface characterization

Electrochemical impedance spectroscopy (EIS) was performed in 1× PBS containing 0.1 M KCl using a CH Instruments 700F bipotentiostat in a three-electrode configuration, with the patterned PBI-LIG electrode serving as both the working and counter electrodes and a commercial Ag/AgCl electrode as the reference. Measurements were collected at a DC bias of 0 V with a 10 mV AC perturbation. The impedance spectrum was recorded from 3 MHz to 1 Hz using 12 points per decade, with a 2 s quiet time before each run. Automatic sensitivity scaling was enabled.

#### 3. Cortisol concentration calibration and analytical performance

A 1 mM cortisol stock solution was prepared by dissolving cortisol in absolute ethanol. Working solutions were generated by serial dilution of the stock into 1× PBS containing 0.1 M KCl, with vigorous vortexing at each dilution step to ensure complete dispersion of the hydrophobic analyte. Fresh dilutions were prepared for each experiment.

Prior to analytical measurements, electrodes were conditioned by five consecutive square-wave voltammetry scans conducted from 0 to −2.0 V using a 10 mV step potential, 250 mV pulse amplitude, and 50 Hz frequency in 1× PBS containing 0.1 M KCl, performed consecutively without inter-scan incubation (**Fig. S15**). After conditioning, the sensor was incubated in blank buffer for 10 min to allow the interfacial response to stabilize, and a blank SWV response was then collected. For subsequent measurements, the sensor was incubated in each test solution for 10 min prior to SWV acquisition. Analytical signals were obtained using the TRI-extracted response, which was plotted against cortisol concentration for calibration and performance analysis.

For each cortisol concentration, the TRI-extracted transition magnitude was recorded. Calibration curves were constructed by plotting the transition magnitude against cortisol concentration. Each concentration was measured in replicate, and the mean ± standard deviation was used for curve fitting. Linear regression was applied over the concentration range in which the response was monotonic and proportional. Analytical sensitivity was defined as the slope of the calibration curve obtained from the TRI-extracted transition magnitude as a function of the logarithm of cortisol concentration. The limit of detection (LOD) was estimated using the 3σ criterion (LOD = 3σ/S), where σ is the standard deviation of replicate blank measurements and S is the slope of the calibration curve.

Selectivity was evaluated by incubating the sensor in solutions containing representative physiological interferents, including progesterone, estradiol, epinephrine, insulin, and norepinephrine. Following a 10 min incubation, SWV measurements were acquired using the standard TRI workflow. Raw derivative-domain responses are shown in **Fig. S12**. Selectivity was quantified by normalizing the TRI-extracted transition magnitude for each interferent to the response obtained for 100 nM cortisol and expressing the result as a percentage.

To assess the temporal response of the sensing interface, devices were incubated in a known cortisol concentration and interrogated after 3, 5, 8, 10, 12, and 15 min incubation periods (Supplementary **Fig. S13**). The response reached a stable profile after approximately 8–10 min; therefore, a 10 min incubation period was used throughout the study. For each time point, an SWV scan was collected following the standard workflow, and the TRI-extracted transition magnitude was used to quantify the time-dependent evolution of the cortisol-linked signal.

Sensors were incubated for 10 min in pH-controlled buffer solutions containing 10 µM cortisol, after which SWV scans were collected. Data were processed using the TRI strategy to extract both the cortisol-associated transition magnitude and the corresponding inflection potential.

To evaluate effects biological ISFs, cortisol standards (1–1000 nM) were prepared in artificial interstitial fluid (aISF). Blank aISF scans were first collected to establish a background signal. Sensors were then incubated in each cortisol-spiked aISF sample for 10 min, scanned using SWV, and the resulting data were background-subtracted using the blank aISF response. TRI-extracted transition magnitudes were used for analytical mapping.

Ex vivo performance was assessed using dermal interstitial fluid extracted from porcine skin. Baseline pig ISF scans were first collected to establish the intrinsic electrochemical response of the native matrix. Sensors were then incubated for 10 min in pig ISF samples spiked with 1 µM or 10 µM cortisol, followed by SWV acquisition. Cortisol-linked responses were extracted and quantified using the TRI analysis framework.

Ex vivo performance was assessed using native human dermal interstitial fluid. Baseline scans were first collected to establish the intrinsic electrochemical response of the native matrix. The same samples were then spiked with 1 µM or 10 µM cortisol, followed by SWV acquisition. Cortisol-linked responses were quantified using the TRI analysis framework.

#### 4. Transition-resolved interrogation (TRI) data analysis

TRI analysis was performed on the raw SWV current–potential responses acquired at each measurement condition. The data were first transformed into the derivative domain (dj/dE). The resulting derivative trace was then processed using the least-squares smoothing function implemented in the CH Instruments software (CHI760F Electrochemical Workstation) with a fixed 49-point smoothing parameter. This smoothing operation employs a Savitzky–Golay least-squares procedure, in which local polynomial fitting is performed across a moving data window to estimate the slowly varying background component. The smoothed derivative trace was subtracted from the original derivative response to generate a residual signal containing localized transition features. The analytical response was defined as the magnitude of the first negative residual lobe and was used for all calibration, selectivity, pH, temporal response, and biological matrix measurements. The same data-processing workflow and smoothing parameters were applied to all datasets throughout the study.

## Data availability

Data that support the findings of this study are available from the corresponding author upon reasonable request.

## Supporting information

not applicable

## Acknowledgment

Z.B. is a Chan Zuckerberg Biohub San Francisco investigator and an Arc Institute innovation investigator. Z.B. acknowledges support from the Chan Zuckerberg Biohub San Francisco, the Tianqiao and Chrissy Chen Ideation and Prototyping Lab and the Stanford Wearable Electronics Initiative (eWEAR) seed funding program. Part of this work was performed at the Stanford Nano Shared Facilities (SNSF), supported by the National Science Foundation under award ECCS-2026822. C.Z. acknowledges support from a fellowship from the National Institute of Biomedical Imaging and Bioengineering of the National Institutes of Health (F32EB034156). C.U. acknowledges support from the Stanford University Medical Scientist Training Program (T32-GM145402). I.W. and K.-J.H. acknowledge support from the Swiss National Science Foundation through postdoctoral fellowships (I.W., P500PT_214498; K.-J.H., P500ON_230529). L.R. acknowledges support from the Niels Stensen Fellowship and the Netherlands Organisation for Scientific Research (NWO) through a Rubicon Fellowship (019.233EN.013). J.H. acknowledges support from the Stanford Cancer Institute, a National Cancer Institute-designated Comprehensive Cancer Center, and the Canary Center at the Stanford School of Medicine. T.W.R. acknowledges support from the NIH Molecular Biophysics Training Program (T32 GM136568). The authors thank Dr. Ena Luis for insightful discussions and feedback on the manuscript.

## Author contributions

A.M. and Z.B. conceived and designed the project. A.M. performed exploratory studies, developed the materials and electrochemical interrogation framework, conducted electrochemical measurements, and carried out data analysis. C.Z. provided experimental assistance and contributed to conceptual discussions, manuscript preparation, and revision. C.U. provided conceptual feedback and assisted with exploratory experiments and manuscript editing. K.-J.H. developed the PBI membrane preprocessing protocol and performed XPS data collection and analysis. I.R.C. performed Raman characterization and contributed to XPS method development. T.C. contributed to XPS data interpretation, assisted with the chemothermal reduction workflow and experimental setup, and contributed to conceptual discussions. B.C.D. and I.J.G. developed the LC–MS/MS method, provided analytical guidance, and collected, processed, and analyzed the LC–MS/MS data. J.L.H., D.I., and T.W.R. provided interstitial fluid samples and related technical support. L.R. contributed to experimental studies and interpretation of surface-chemistry changes associated with chemothermal treatment. S.W. and I.W. contributed expertise in LIG-based electrochemical sensing. D.P. collected SEM images. H.L. contributed to early experimental planning and discussions. Y.C. contributed to discussions on graphene characterization, and C.X. contributed expertise in LIG technology. B.S. designed the flexible interconnects, and A.S. assisted with exploratory experiments. J.H. supported materials characterization. J.M.D. supervised the interstitial fluid sampling collaboration. A.M. drafted the manuscript. Z.B. and C.Z. contributed to manuscript revision and editing. Z.B. supervised the project. All authors approved the final manuscript.

