## Supplementary material for "Direct electrochemical cortisol detection via transition-resolved interrogation at a defect-engineered graphene interface": not applicable

**Supplementary Note 1 — Electrochemical behavior of cortisol in aprotic media**

Cortisol exhibits weak intrinsic electrochemical activity in aprotic media under conventional electrochemical interrogation. Classical studies of corticosteroids report broad, low-intensity reduction waves at relatively large overpotentials, consistent with kinetically limited, irreversible electron transfer (1). More broadly, the limited electroactivity of steroid carbonyls has been attributed to poor stabilization of reduced intermediates and the substantial driving force required for electron uptake (2).

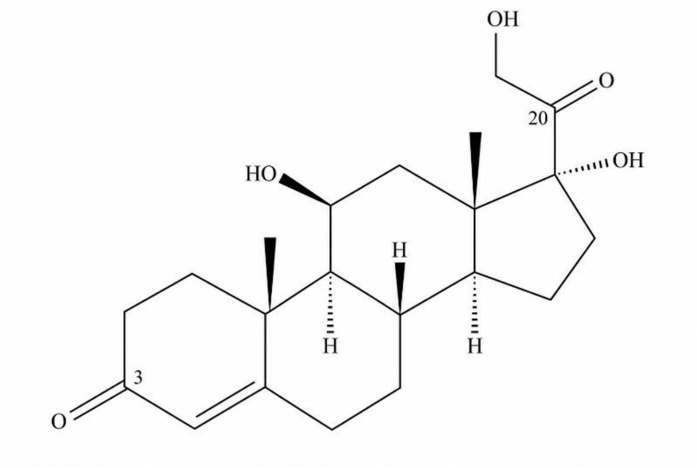


**Supplementary Fig. S1. Molecular structure of cortisol.** The C3 and C20 carbonyl groups are the most plausible sites for electrochemical reduction. This assignment is consistent with the weak, poorly resolved reductive responses reported for corticosteroids under conventional voltammetric interrogation.

To determine whether cortisol exhibits measurable electroactivity under well-controlled conditions, we examined its behavior in an aprotic environment using cyclic voltammetry (CV) and square-wave voltammetry (SWV). Measurements were performed using a glassy carbon working electrode, a platinum counter electrode, and an $Ag/Ag^{+}$ reference electrode in dry acetonitrile (MeCN) containing 0.1 M tetrabutylammonium hexafluorophosphate ($TBAPF_6$). Ferrocene/ferrocenium ($Fc/Fc^{+}$) was included as an internal standard, and all potentials are reported versus $Fc/Fc^{+}$.

### Cyclic Voltammetry

Cyclic voltammetry showed that cortisol is measurably electroactive in aprotic media, but only at high negative cathodic potentials (Fig. S2). The broad cathodic feature and weak anodic response on the reverse sweep indicate quasi-reversible electrochemical behavior.

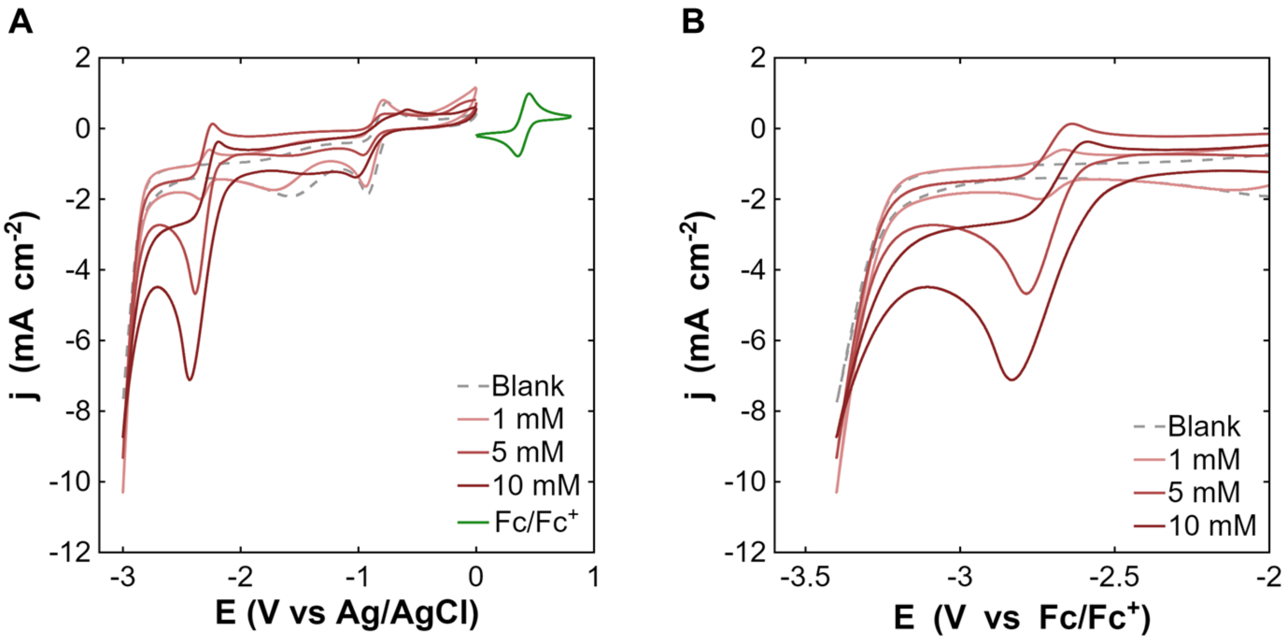


**Supplementary Fig. S2. Cyclic voltammetry of cortisol in dry MeCN.** (A) Full cyclic voltammograms recorded in dry MeCN containing 0.1 M $TBAPF_6$, showing the concentration-dependent cortisol reduction response together with the 1 mM $Fc/Fc^+$ redox couple used as an internal reference. Background features centered at approximately −0.9 and −1.6 V are also observed in the blank electrolyte and remain distinct from the cortisol-associated reduction at more negative potentials. (B) Expanded view of the cortisol reduction region. Cortisol exhibits a broad cathodic peak centered at approximately −2.8 V versus $Fc/Fc^+$ that is absent in the blank electrolyte and increases systematically with cortisol concentration, confirming its faradaic origin. A weak anodic feature appears on the reverse sweep, indicating limited reversibility under these measurement conditions.

**Square-Wave Voltammetry**

Square-wave voltammetry provided enhanced sensitivity to cortisol's faradaic response, revealing a concentration-dependent cathodic feature centered at approximately −2.45 V versus $Fc/Fc^+$ (Fig. S3). Similar to cyclic voltammetry, SWV also resolved a measurable reverse component, consistent with partial re-oxidation of the reduced species on the shorter pulse timescale. Together, these data confirm that cortisol exhibits measurable intrinsic electroactivity under controlled aprotic conditions.

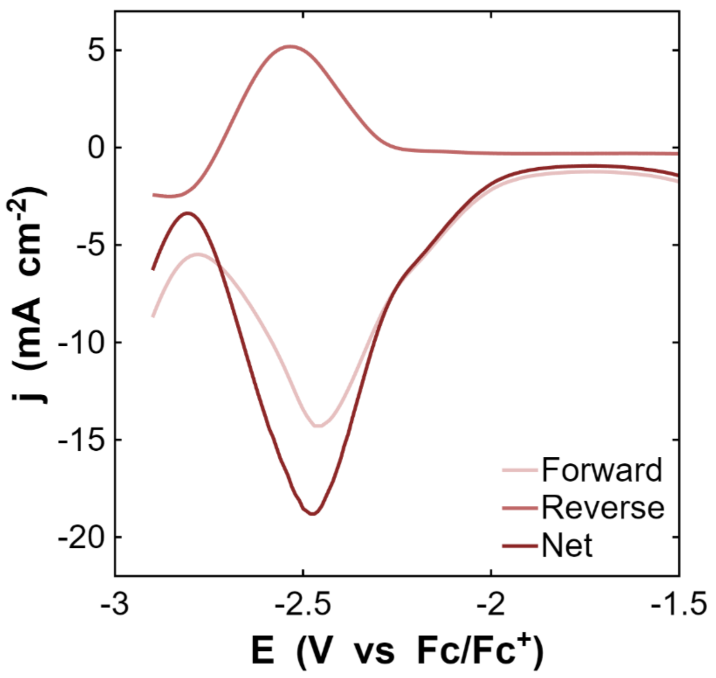


**Supplementary Fig. S3. Square-wave voltammetry of cortisol in dry MeCN.** Rrepresentative forward, reverse, and net square-wave voltammetric responses for cortisol recorded on a glassy carbon working electrode in dry MeCN containing 0.1 M $TBAPF_6$. A concentration-dependent cathodic response is observed in the forward scan, accompanied by a measurable anodic reverse component consistent with partial re-oxidation of the reduced species. The resulting net response confirms the faradaic origin of the cortisol signal under pulsed interrogation conditions.

**Supplementary Note 2 — Electrochemical reduction of cortisol in aqueous media**

Cortisol's electrochemical behavior changes substantially in aqueous electrolyte, where proton availability and competing cathodic processes alter the observed reduction response. We therefore examined its behavior in a controlled aqueous electrolyte consisting of 1× PBS containing 0.1 M KCl. All potentials in this Note are reported versus Ag/AgCl.

**Cyclic Voltammetry**

Cyclic voltammetry revealed a broad cathodic feature centered near −1.75 V versus Ag/AgCl that was absent in the blank electrolyte (Fig. S4), confirming measurable electrochemical activity in aqueous media. However, this response overlaps substantially with the onset of hydrogen evolution, producing merged cathodic currents that obscure the cortisol-associated feature and preclude direct quantification using conventional voltammetry.

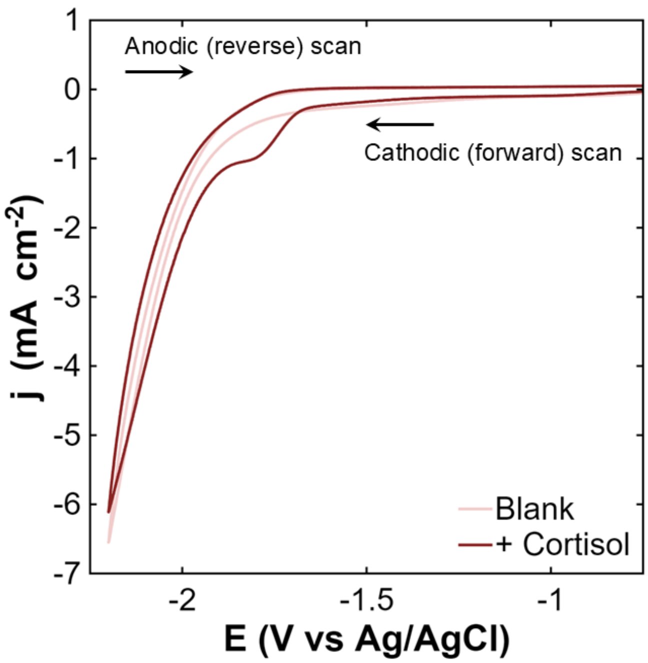


**Supplementary Fig. S4. Cyclic voltammograms of cortisol in aqueous buffer.** Representative cyclic voltammograms recorded at 0.5 V $s^{-1}$ on a glassy carbon working electrode in 1× PBS containing 0.1 M KCl. A broad cathodic feature centered near –1.75 V vs Ag/AgCl is observed for 10 mM cortisol and is absent in the blank electrolyte, confirming measurable electrochemical activity in aqueous media. Substantial overlap with the onset of hydrogen evolution produces merged cathodic currents that obscure direct resolution of the cortisol-associated feature under conventional voltammetric interrogation.

**Square Wave Voltammetry**

Square-wave voltammetry in PBS similarly revealed a weak, poorly resolved cathodic response with no clearly discernible reverse component. The overlap with hydrogen evolution complicates direct signal assignment and underscores the challenge of selectively resolving the cortisol-associated transition using conventional electrochemical interrogation.

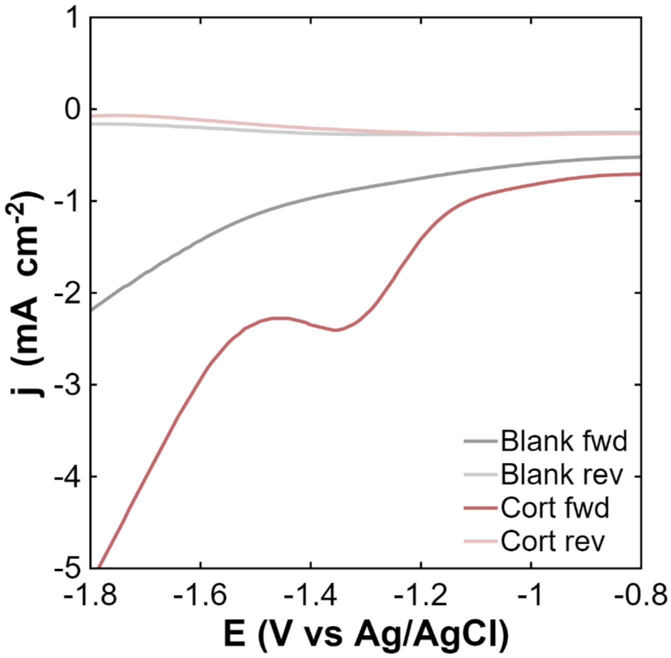


**Supplementary Fig. S5. Square-wave voltammetry of cortisol in aqueous buffer.** Representative forward and reverse square-wave voltammetric responses recorded on a glassy carbon working electrode in 1× PBS containing 0.1 M KCl for the blank electrolyte and for 10 mM cortisol. The cortisol trace exhibits a broad cathodic feature that appears predominantly in the forward pulse, with no well-defined corresponding reverse signal, consistent with kinetically limited reduction behavior under aqueous conditions. This feature emerges within the same potential region as the onset of hydrogen evolution, resulting in overlapping cathodic currents that hinder direct resolution of the cortisol-associated response under conventional interrogation conditions.

**Supplementary Note 3 — Physical basis of transition-resolved interrogation**

Transition-resolved interrogation (TRI) is a waveform-based electrochemical interrogation regime we designed to resolve electrochemical processes that evolve on distinct timescales under competitive cathodic conditions. TRI uses a square-wave waveform superimposed on a staircase potential sweep, similar in structural form to square-wave voltammetry (3). Although square-wave voltammetry is often associated with reversible redox analysis, its timing architecture can also provide useful kinetic discrimination for irreversible electrochemical processes (4). We design TRI to operate in a distinct regime in which waveform timing and amplitude are used to selectively control which competing interfacial processes are allowed to develop prior to measurement.

**3.1 Time-domain competition between interfacial processes**

During TRI, the measured current arises from the superposition of multiple interfacial processes:

$$I(t,E) = I_C(t,E) + I_{cort}(t,E) + I_{HER}(t,E) + \epsilon$$

where $I_C$ represents capacitive charging, $I_{cort}$ the cortisol-associated faradaic response, $I_{HER}$ the competing hydrogen evolution current, and $\epsilon$ bounded measurement noise.

These processes evolve on distinct characteristic timescales. Capacitive charging is a rapid interfacial process associated with charging and discharging of the electrochemical double layer. The resulting non-Faradaic current dominates immediately following a potential transition but decays rapidly as the double layer equilibrates (5). In contrast, the cortisol-associated faradaic current develops over a finite timescale under cathodic polarization, becoming observable only after sufficient overpotential is established and the associated redox process is activated. Hydrogen evolution emerges more slowly, requiring sufficient overpotential and time for nucleation and growth of hydrogen bubbles associated with the competing cathodic reaction (6). These distinct temporal behaviors provide the basis for TRI.

**3.2 Waveform timing as the control parameter**

For a square-wave waveform applied at frequency $f$, the effective dwell time at each polarization state is

$$\tau_d = \frac{1}{2f}$$

which defines the time available for interfacial processes to evolve before current sampling.

We choose a regime where the dwell time exceeds the rapid decay of capacitive charging but remains shorter than the timescale required for substantial hydrogen evolution:

$$\tau_C \ll \tau_d \ll \tau_{HER}$$

where $\tau_C$ represents the characteristic capacitive relaxation timescale and $\tau_{HER}$ the effective onset timescale of hydrogen evolution.

Under these conditions, the cortisol-associated transition becomes observable before hydrogen evolution dominates the cathodic response. The waveform therefore acts as a temporal gate, favoring current from the analyte-linked Faradaic response over competing background processes.

This temporal dependence defines three operating regimes:

- **Short dwell times (high frequency):** residual capacitive charging dominates, limiting analyte visibility.
- **Long dwell times (low frequency):** hydrogen evolution develops substantially and dominates the cathodic response.
- **Intermediate dwell times:** the cortisol-associated response becomes observable while hydrogen evolution and capacitive current are both minimized, defining the operative TRI window.

#### 3.3 Mapping temporal kinetics into the sampled current–potential response

Because current is sampled after a fixed dwell interval at each potential step, temporal differences between competing interfacial processes are encoded into the measured current–potential response.

Each sampled point therefore reflects the state of the interface after evolving for a defined interval under a specific polarization condition, rather than an instantaneous equilibrium response. In this way, waveform timing converts time-domain kinetic competition into a measurable potential-domain signal structure. Under the intermediate dwell-time regime, the cortisol-associated reduction emerges as a localized feature embedded within a broader evolving cathodic background. The appearance and prominence of this feature are therefore determined not only by the intrinsic electrochemistry of cortisol, but by the waveform-imposed measurement conditions.

TRI utilizes the timing-based discrimination of SWV to distinguish the interfacial kinetic landscape during signal acquisition. Instead of relying on forward–reverse subtraction, TRI uses a frequency-tuned, high-amplitude square-wave excitation to impose a controlled dwell time at each polarization state. This dwell time determines the extent to which competing interfacial processes contribute to the measured overall signal.

#### 3.4 Frequency-controlled kinetic selectivity

The degree of kinetic discrimination achieved by TRI depends directly on waveform timing. Experimentally, this behavior is quantified using the selectivity metric

$$S(f) = \frac{A_{cort}(f)}{A_{HER}(f)}$$

where $A_{cort}$ represents the extracted amplitude of the cortisol-associated transition and $A_{HER}$ the competing hydrogen evolution contribution at excitation frequency $f$.

Maximization of $S(f)$ identifies the operating regime in which analyte observability is favored relative to competing cathodic background processes.

#### 3.5 First-derivative representation and background estimation

To improve observability of the cortisol-associated feature, the measured current–potential response is represented in first-derivative form:

$$z(E) = \frac{dI_m}{dE}$$

where $I_m(E)$ is the sampled current response.

This transformation suppresses slowly varying background contributions while emphasizing more rapidly evolving localized changes associated with the cortisol reduction. Because hydrogen evolution and baseline drift evolve gradually across the cathodic potential window, whereas the cortisol-associated response emerges over a more confined activation region, derivative representation improves separation between the analyte-linked feature and the broader background.

To further isolate the transition, the slowly varying background is estimated using least-squares smoothing:

$$\hat{z}_{base}(E) = S\{z(E)\}$$

where $S$ denotes the smoothing operator. The transition-resolved component is then obtained by subtraction:

$$z_{cort}(E) = z(E) - \hat{z}_{base}(E)$$

Smoothing-based baseline estimation and subtraction have been widely employed for background correction and feature extraction in analytical signal processing (7,8). This procedure suppresses the slowly varying background while preserving localized features associated with the cortisol-linked transition, thereby improving contrast and facilitating quantitative analysis.

**Supplementary Note 4 — Benchmarking polybenzimidazole-derived LIG (PBI-LIG) against polyimide-derived LIG (PI-LIG)**

Polyimide-derived laser-induced graphene (PI-LIG) is a widely used benchmark material in electrochemical sensing (9–11). To contextualize the properties of the polybenzimidazole-derived laser-induced graphene (PBI-LIG) interface used in this work, we compared its structural, compositional, and electrochemical characteristics against PI-LIG.

Laser-induced carbonization of PBI produces a materially distinct carbon architecture relative to PI-LIG. Raman spectroscopy (Supplementary Fig. 6A) shows characteristic D and G bands for both materials, with PBI-LIG exhibiting a higher D/G intensity ratio (2.03 vs 1.23), consistent with increased defect density and heteroatom incorporation. X-ray photoelectron spectroscopy (Supplementary Fig. 6B) further highlights compositional differences, with PBI-LIG containing substantially higher nitrogen (8.18 at% vs 2.7 at%) and oxygen content (18.3 at% vs 9.52 at%). A feature at ~505 eV corresponds to the Na KLL Auger transition, likely arising from trace sodium retained during PBI membrane processing.

Electrochemical characterization revealed consistent performance advantages for PBI-LIG. Cyclic voltammetry in PBS (Supplementary Fig. 6C) showed a larger capacitive envelope, consistent with increased electrochemically accessible surface area. Electrochemical impedance spectroscopy (Supplementary Fig. 6D) showed lower charge-transfer resistance for PBI-LIG. Ferri/ferrocyanide redox probing (Supplementary Fig. 6E) further indicated faster heterogeneous electron-transfer kinetics, with PBI-LIG approaching near-Nernstian peak separations (≈56–62 mV) compared with substantially larger separations for PI-LIG (≈92–103 mV). Together, these data establish PBI-LIG as a more electrochemically active interface than the conventional PI-LIG benchmark.

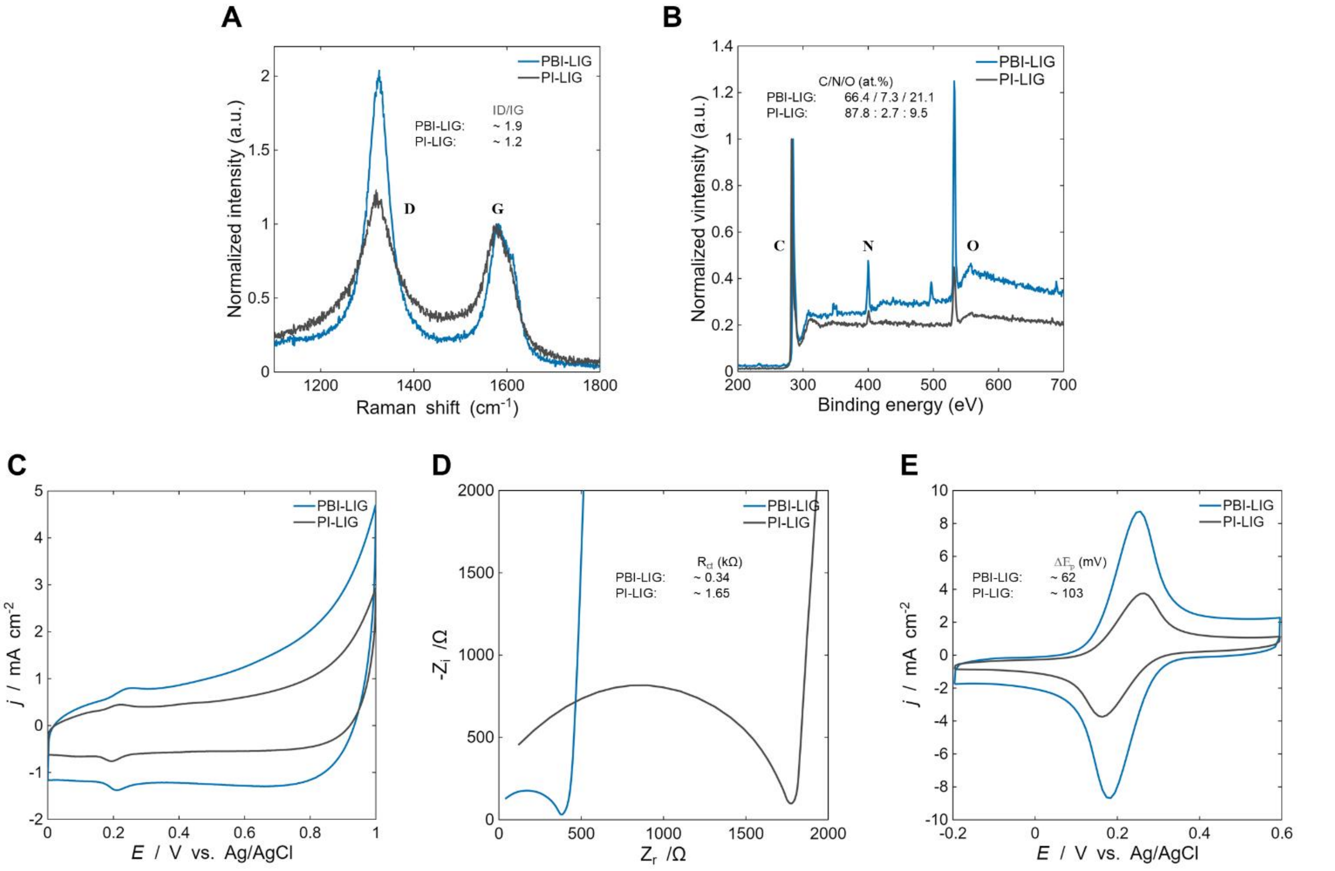


**Supplementary Fig. S6. Structural, compositional, and electrochemical comparison of PBI-LIG and PI-LIG.** (A) Raman spectra showing the characteristic D and G bands for both materials. PBI-LIG exhibits a higher D/G intensity ratio, consistent with increased defect density and heteroatom incorporation relative to PI-LIG. (B) X-ray photoelectron spectroscopy (XPS) survey spectra showing higher nitrogen and oxygen content in PBI-LIG relative to PI-LIG, consistent with compositional differences arising from precursor chemistry and laser-induced carbonization. (C) Cyclic voltammetry in PBS within the non-Faradaic region, showing a larger capacitive envelope for PBI-LIG, indicative of increased electrochemically accessible surface area. (D) Electrochemical impedance spectroscopy (Nyquist plots) demonstrating lower charge-transfer resistance for PBI-LIG. (E) Ferri/ferrocyanide redox probing showing higher peak currents and smaller peak-to-peak separation ($\Delta E_p$) for PBI-LIG, consistent with faster heterogeneous electron-transfer kinetics.

**Supplementary Note 5 — Chemothermal modulation of PBI-LIG**

The chemothermal treatment consists of a two-step reduction and annealing process. PBI-LIG electrodes were first treated with hydrazine monohydrate at 95 °C under nitrogen for 24 h, followed by a mild vacuum annealing step at 100 °C for 20 h. Hydrazine reduction targets oxygen-containing surface functionalities, whereas the subsequent annealing step removes residual adsorbates and weakly bound surface species, yielding a cleaner graphene interface for electrochemical interrogation. To assess the effects of chemothermal treatment on the electrochemical interface, we compared as-printed and hydrazine-treated/annealed PBI-LIG using structural, compositional, and electrochemical characterization (Supplementary Fig. 7).

Raman spectra retained the characteristic D, G, and 2D bands following treatment, with D/G decreasing from ~1.9 to ~1.4 and 2D/G increasing from ~0.25 to ~0.31, indicating preservation of the graphitic framework with partial restoration of $sp^2$ ordering. Square-wave voltammetry in blank buffer reveals distinct changes in the electrochemical response after treatment, indicating modification of surface-accessible redox states. X-ray photoelectron spectroscopy of the N 1s region revealed substantial redistribution of nitrogen chemical environments following chemothermal processing (Fig. S7). While the total nitrogen content decreased modestly from 7.3 to 6.6 at.%, deconvolution of the N 1s envelope showed that the relative pyridinic N contribution increased from ~14% to ~37%, while the pyrrolic N contribution decreased from ~67% to ~42%; the graphitic N fraction remained comparatively unchanged (~19–20%). Together with the concurrent reduction in oxygen content, these observations indicate substantial redistribution of the surface nitrogen chemical environments and modification of the local interfacial chemistry of PBI-LIG following chemothermal treatment.

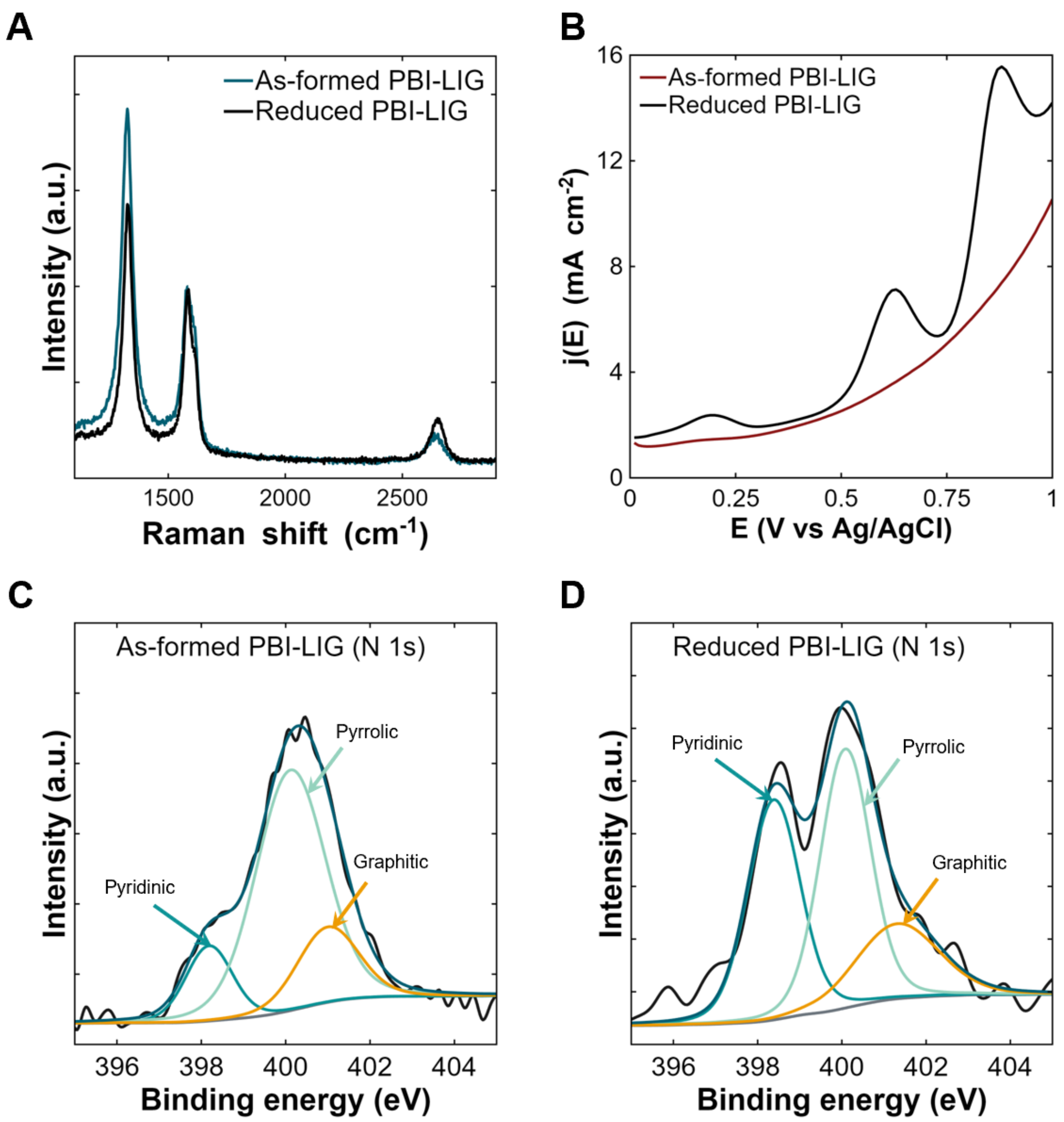


**Supplementary Fig. S7. Chemothermal modulation of PBI-LIG.** (A) Raman spectra of as-printed and chemothermally treated PBI-LIG, showing preservation of the graphitic carbon framework with modest changes in defect-associated spectral features following treatment. (B) Square-wave voltammetry (0–1 V) in blank buffer showing altered electrochemical response after chemothermal treatment, consistent with modification of surface-accessible redox states. (C) N 1s XPS spectrum of as-printed PBI-LIG showing pyridinic, pyrrolic, and graphitic nitrogen components. (D) N 1s XPS spectrum of chemothermally treated PBI-LIG showing redistribution of nitrogen chemical environments following processing.

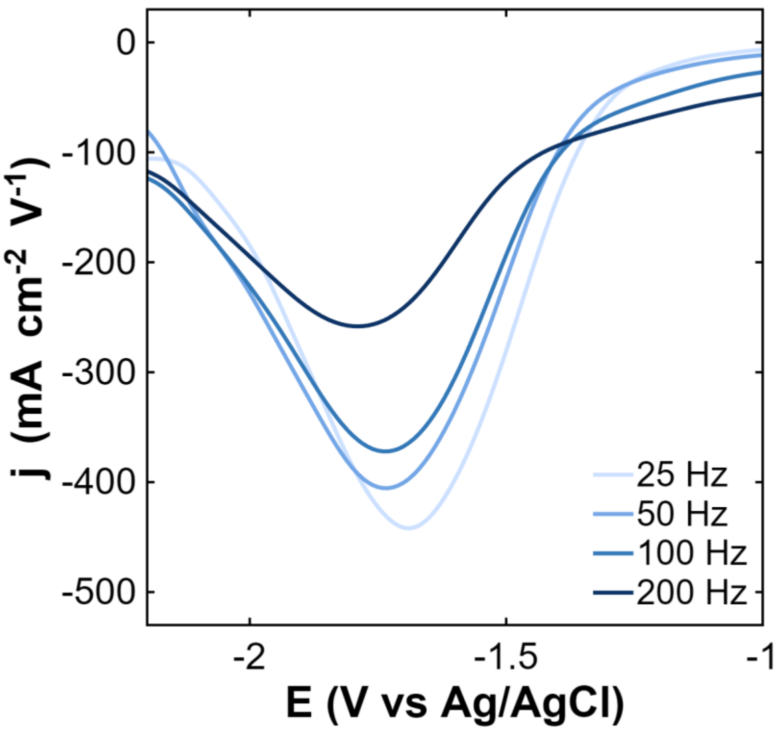


**Supplementary Fig. S8. Frequency-dependent hydrogen evolution on reduced PBI-LIG.** Square-wave voltammograms collected on reduced PBI-LIG in 1× PBS + 0.1 M KCl showing the hydrogen-evolution response at 25, 50, 100, and 200 Hz. The HER-only signal is substantially larger at lower frequencies, consistent with longer dwell times allowing nucleation and growth processes to develop. At higher frequencies, the shortened dwell time suppresses HER, yielding a markedly reduced faradaic contribution. These measurements demonstrate that longer dwell times promote HER development, whereas higher-frequency interrogation kinetically limits HER on reduced PBI-LIG.

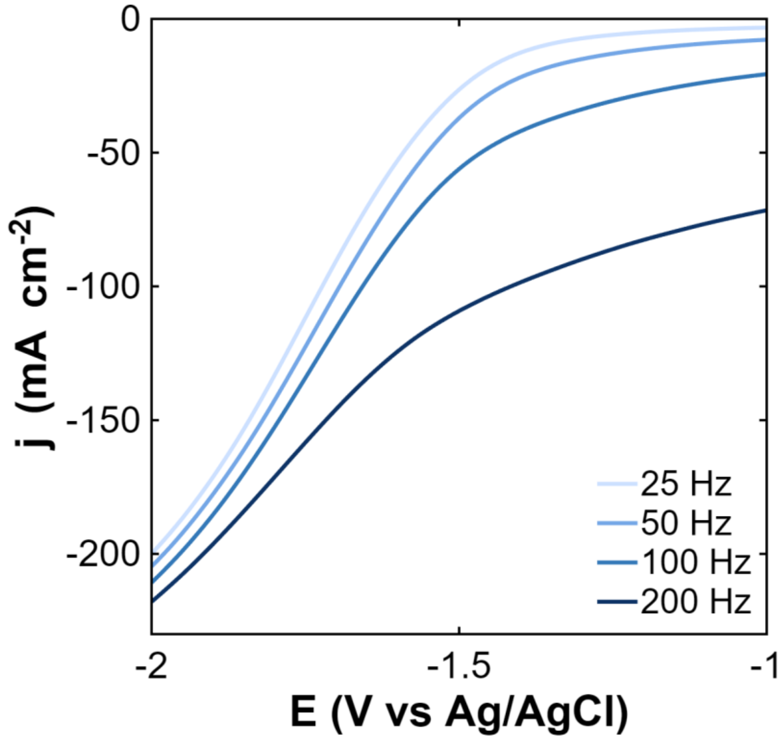


**Supplementary Fig. S9. Frequency-dependent capacitive background on reduced PBI-LIG in the presence of cortisol.** Raw square-wave voltammograms collected on reduced PBI-LIG in 1× PBS + 0.1 M KCl with 100 µM cortisol at 25, 50, 100, and 200 Hz. The non-faradaic background increases substantially at higher frequencies, reflecting the larger capacitive charging currents associated with very short dwell times. As the dwell time decreases, the capacitive component dominates the total current, obscuring weak faradaic features and supporting the conclusion that short-dwell-time interrogation significantly amplifies the non-faradaic background.

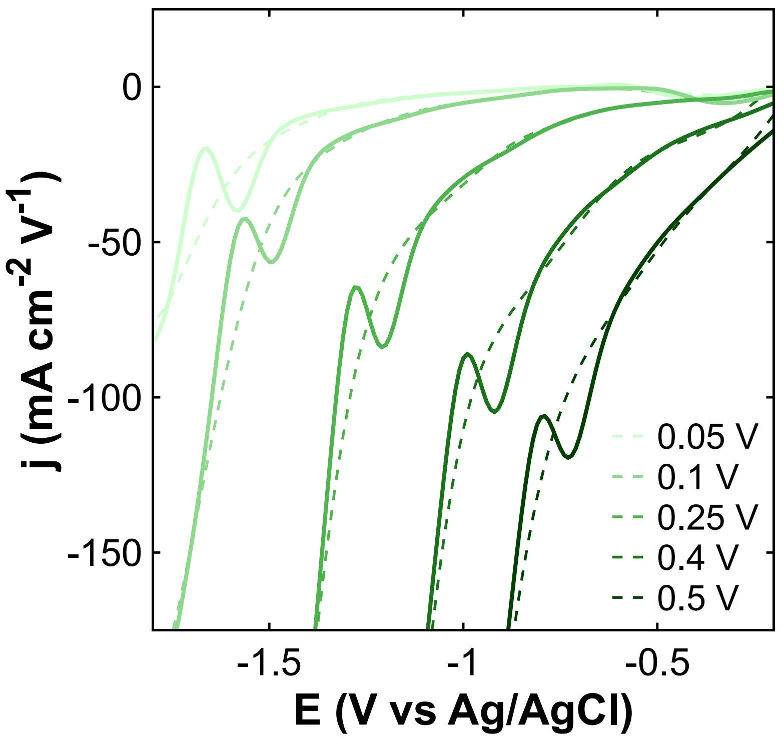


**Supplementary Fig. S10. Amplitude-dependent derivative-domain SWV transition responses (dj/dE vs E).** Derivative-domain SWV transition responses collected at excitation amplitudes of 0.05, 0.1, 0.25, 0.4, and 0.5 V using 100 µM cortisol in 1× PBS containing 0.1 M KCl. Solid lines show the cortisol-containing measurements, and dashed lines show the corresponding background responses at each excitation amplitude. Measurements were performed on reduced PBI-LIG electrodes using a frequency of 50 Hz and a step potential of 10 mV. The apparent cortisol-linked transition potential shifts monotonically toward more positive values with increasing excitation amplitude. Low amplitudes yield weak transition features, whereas high amplitudes introduce substantial background growth that reduces analyte-to-background contrast. These data illustrate how excitation amplitude governs both the apparent transition potential and the achievable analyte-to-background contrast.

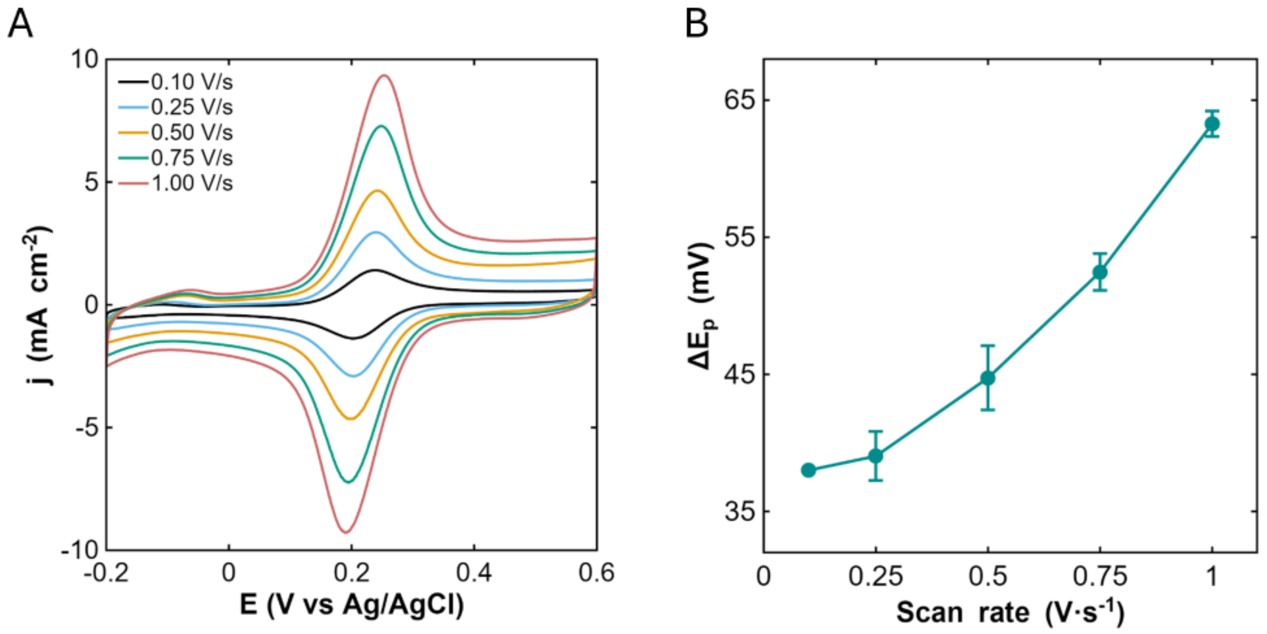


**Supplementary Fig. S11. Ferri/ferrocyanide electron-transfer characteristics of laser-written PBI-LIG.** (A) Cyclic voltammograms of 2 mM ferri/ferrocyanide in 1× PBS containing 0.1 M KCl collected at scan rates from 0.1 to 1.0 V $s^{-1}$. (B) Peak-to-peak separation (ΔEp) extracted from the voltammograms in (A) plotted as a function of scan rate. The relatively small ΔEp values (≈38–63 mV) across the investigated scan-rate range are consistent with efficient heterogeneous electron transfer at the defect-rich PBI-LIG interface.

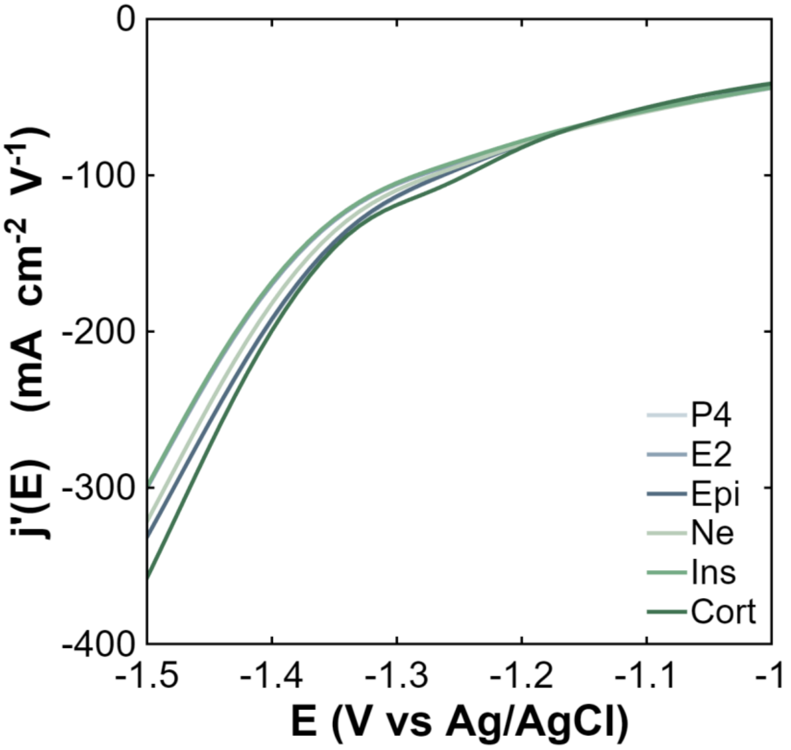


**Supplementary Fig. S12. Raw derivative-domain responses underlying the selectivity analysis in Fig. 4B**. Measurements were acquired under identical TRI conditions following 10 min incubation in PBS containing 100 nM cortisol (Cort) or representative physiological interferents (200 pM progesterone (P4), 10 nM estradiol (E2), 10 nM epinephrine (Epi), 10 nM norepinephrine (Ne), and 1 nM insulin (Ins)).

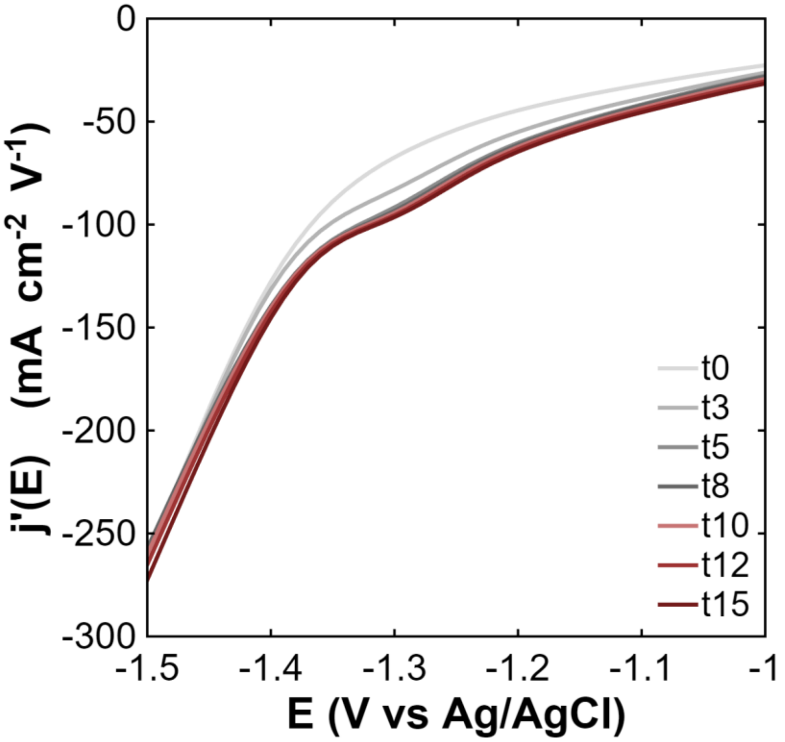


**Supplementary Fig. S13. Temporal evolution of the electrochemical response following cortisol exposure.** Measurements were collected following incubation of the sensor in 100 nM cortisol for 0, 3, 5, 8, 10, 12, and 15 min. The response progressively evolves during the initial incubation period and reaches a stable profile after approximately 8–10 min. The close overlap of the responses acquired at 10, 12, and 15 min indicates that the sensing interface has reached equilibrium under the measurement conditions used. Consequently, a 10 min incubation period was employed throughout the study to ensure measurements were acquired after equilibration.

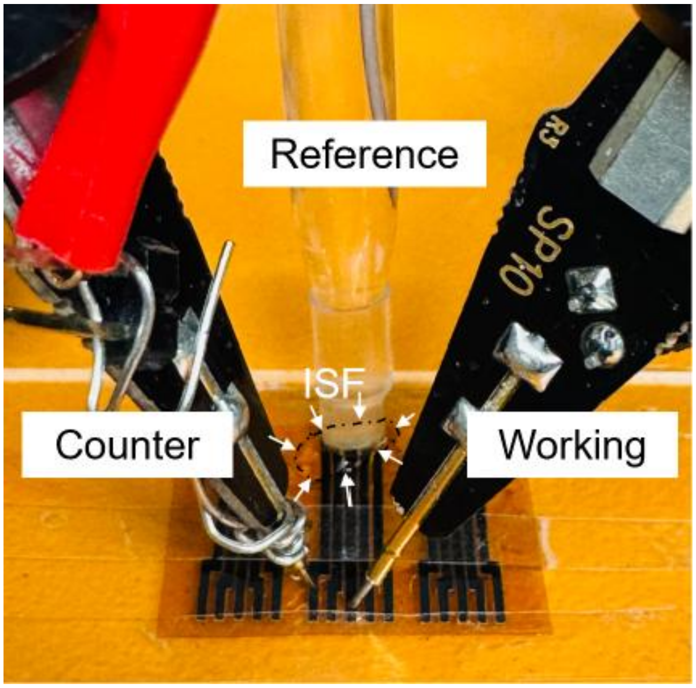


**Supplementary Fig. S14. Electrochemical measurement setup and implantable device architecture.** Representative photograph of the three-electrode configuration used for ex vivo ISF measurements, comprising reduced PBI-LIG working and counter electrodes and a commercial Ag/AgCl reference electrode. The photograph shows three copies of the implantable device; each device contains three working electrodes, one counter electrode, and an additional carbon electrode designed for modification with Ag/AgCl paste to serve as an integrated reference electrode during *in vivo* measurements. White arrows indicate the location of the ISF sample contacting the measurement electrodes. Approximately 10–15 µL of ISF was used per measurement.

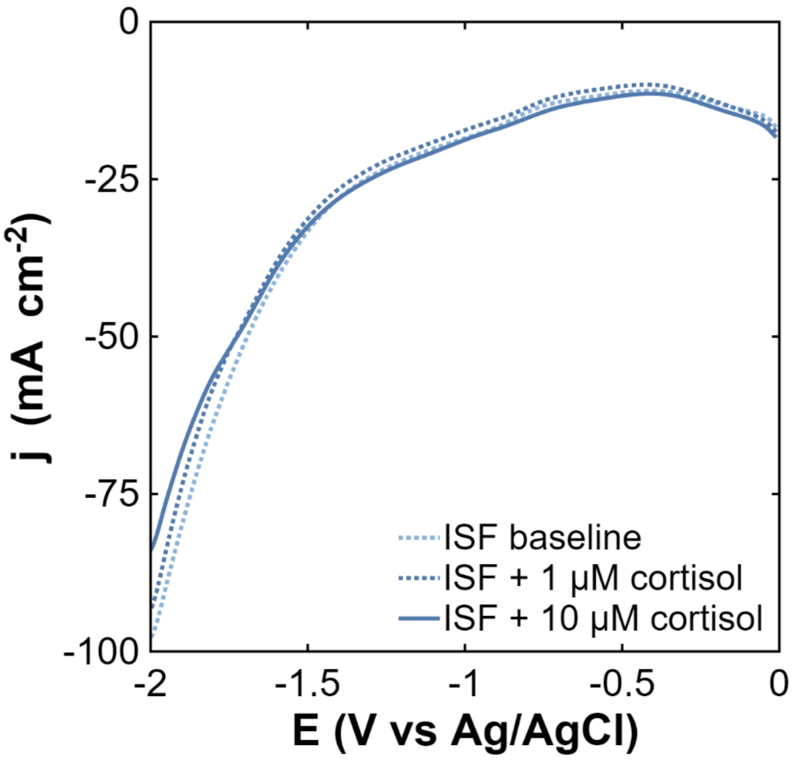


**Supplementary Fig. S15. Raw electrochemical responses in porcine dermal interstitial fluid.** Representative raw SWV j–V curves measured in native porcine dermal ISF and after 10 min incubation in porcine ISF spiked with 1 or 10 μM cortisol. The cortisol-linked response is embedded within the broader matrix background and resolved through TRI analysis in **Fig. 5d**. Measurements were independently performed using three sensors; representative data from one sensor are shown.

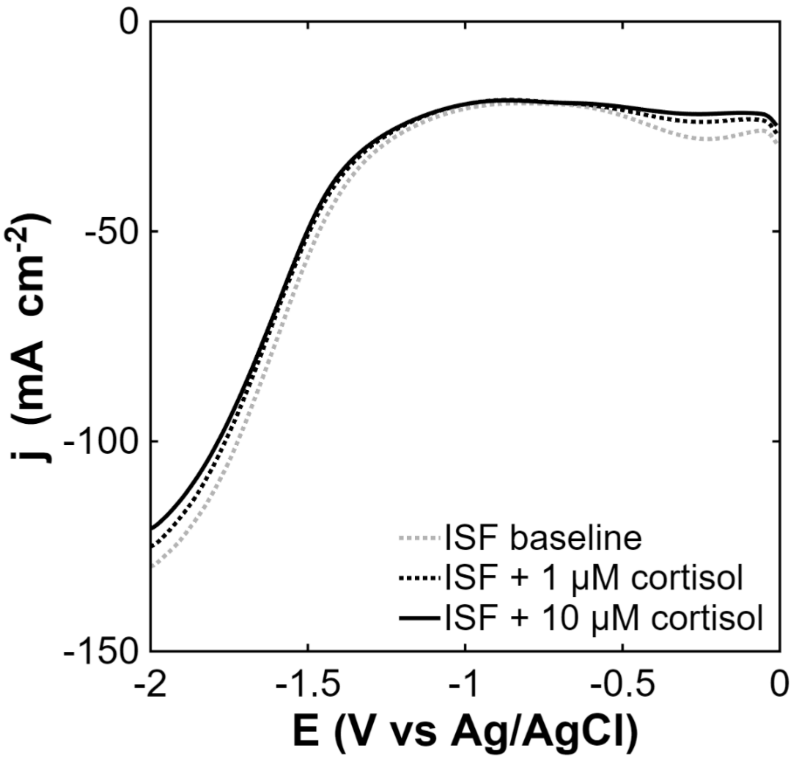


**Supplementary Fig. S16. Raw electrochemical responses in native human dermal interstitial fluid.** Representative raw square-wave voltammetry (SWV) j–V curves measured using a defect-engineered graphene sensor in native human dermal ISF before and after addition of 1 or 10 µM cortisol. Measurements were acquired from 0 to −2.0 V versus Ag/AgCl using a 10 mV step potential, 250 mV pulse amplitude, and 50 Hz frequency. The cortisol-linked faradaic response is embedded within the broader matrix background in the raw voltammograms and is resolved through TRI analysis in **Fig. 5e**. Measurements were independently performed using three sensors; representative data from one sensor are shown.

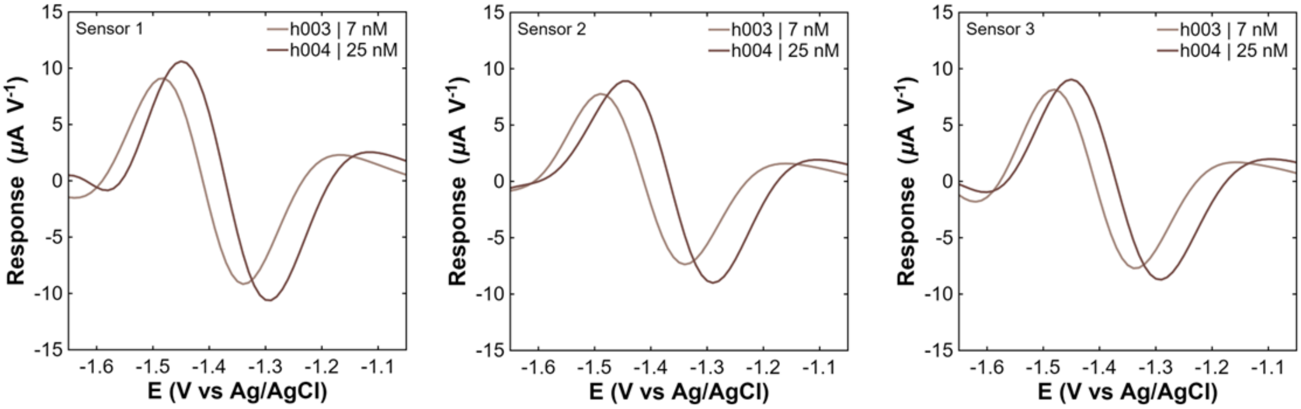


**Supplementary Fig. S17. Reproducibility of endogenous cortisol detection in native human ISF.** Extracted TRI signatures measured using three independent sensors in native human ISF samples h003 and h004, containing 7 and 25 nM cortisol, respectively, as independently determined by LC–MS/MS. Each panel shows the individual responses obtained with one sensor. All three sensors resolved the cortisol-linked transition, with the higher-concentration h004 sample producing a larger transition magnitude and a slight shift in apparent reduction potential relative to h003. These potential differences may arise from variations in pH, ionic strength, or other matrix constituents between the samples.

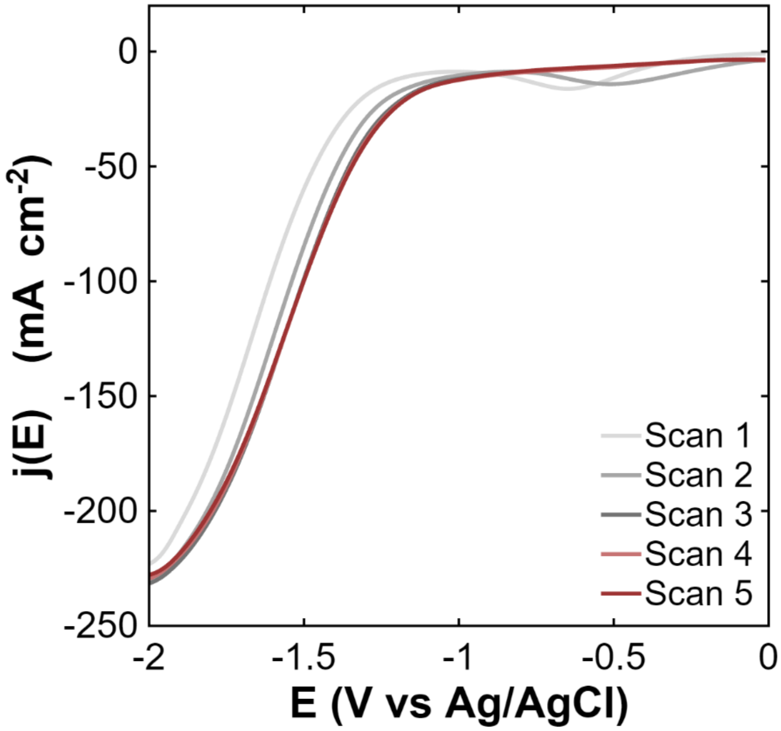


**Supplementary Fig. S18. Electrochemical conditioning of the defect-engineered graphene interface.** Five consecutive square-wave voltammetry (SWV) conditioning scans collected on the defect-engineered graphene interface in 1× PBS containing 0.1 M KCl. Scans were performed consecutively without inter-scan incubation using a 10 mV step potential, 250 mV pulse amplitude, and 50 Hz frequency. The first two scans exhibit a pronounced cathodic feature that progressively diminishes with repeated interrogation, consistent with electrochemical reduction of an initially accessible interfacial species. By the third scan, the voltammetric response reaches a stable profile, and subsequent scans (Scans 3–5) are highly reproducible, indicating that the interface has reached a steady electrochemical state. This conditioning protocol establishes a stable electrochemical baseline prior to TRI-based cortisol interrogation.
